# Cryo-EM Unveils the Processivity Mechanism of Kinesin KIF1A and the Impact of its Pathogenic Variant P305L

**DOI:** 10.1101/2023.02.02.526913

**Authors:** Matthieu P.M.H. Benoit, Lu Rao, Ana B. Asenjo, Arne Gennerich, Hernando Sosa

## Abstract

Mutations in the microtubule-associated motor protein KIF1A lead to severe neurological conditions known as KIF1A-associated neurological disorders (KAND). Despite insights into its molecular mechanism, high-resolution structures of KIF1A-microtubule complexes remain undefined. Here, we present 2.7-3.4 Å resolution structures of dimeric microtubule-bound KIF1A, including the pathogenic P305L mutant, across various nucleotide states. Our structures reveal that KIF1A binds microtubules in one- and two-heads-bound configurations, with both heads exhibiting distinct conformations with tight inter-head connection. Notably, KIF1A’s class-specific loop 12 (K-loop) forms electrostatic interactions with the C-terminal tails of both α- and β-tubulin. The P305L mutation does not disrupt these interactions but alters loop-12’s conformation, impairing strong microtubule-binding. Structure-function analysis reveals the K-loop and head-head coordination as major determinants of KIF1A’s superprocessive motility. Our findings advance the understanding of KIF1A’s molecular mechanism and provide a basis for developing structure-guided therapeutics against KAND.

## Introduction

KIF1A, a neuron-specific member of the kinesin-3 family, is a microtubule (MT) plus-end-directed motor proteins that plays a key role in the migration of nuclei in differentiating brain stem cells^1,2^ and the transport of synaptic precursors and dense core vesicles to axon terminals^3–8^. The dysfunction of KIF1A is linked to a spectrum of severe neurodevelopmental and neurodegenerative diseases known as KIF1A-associated neurological disorders (KAND). These disorders include progressive spastic paraplegias, microcephaly, encephalopathies, intellectual disability, autism, autonomic and peripheral neuropathy, optic nerve atrophy, cerebral and cerebellar atrophy, and others^9–51^. To date, more than 145 inherited and *de novo* KAND mutations have been identified, and these mutations span the entirety of the KIF1A protein sequence^9^. The majority are located within the motor domain (MD or ‘head’)^9^ and are thus predicted to affect the motor’s motility properties whereas mutations located outside the motor domain are likely involved in mediating dimerization, autoinhibition, and/or cargo binding^10^. Our own work^9,52^ and the work of others^53,54^ has indeed shown that KAND motor domain mutations affect the motor’s ability to generate force and movement.

Through the patient advocacy group KIF1A.org, more than 580 families with children and adults with KIF1A mutations have come together to improve the lives of those affected by KAND and to accelerate drug discovery. Despite these efforts, the molecular etiologies of KAND remain poorly understood, in part because KIF1A’s molecular mechanism remains unclear. For example, KIF1A is extremely fast and super-processive^55–59^ (the motor can take more than a thousand steps before dissociating), and at the same time, gives up easily under load^52^. These behaviors distinguish KIF1A from the founding member of the kinesin family, kinesin-1^60,61^, and it remains unknown how KIF1A achieves this unique set of properties.

While the motor domain of KIF1A is highly conserved among the members of the kinesin superfamily^62,63^, it possesses distinct structural elements that may account for KIF1A’s unique set of properties. Notably, KIF1A’s positively charged loop 12, known as the “K-loop” due to its KNKKKKK sequence, facilitates the initial binding of KIF1A to the MT^64^ and is crucial for the motor’s rapid reattachment to the MT following detachment under load^52^. This reattachment ability results in a unique sawtooth-like pattern in force generation events^52^. Despite these insights, our understanding of KIF1A’s unique motile properties remains limited due to the absence of high-resolution structural data of the KIF1A-MT complex. The KIF1A motor domain has been visualized under various experimental conditions using X-ray crystallography^65–68^ and in complex with MTs at medium resolution (6-15 Å) using cryoEM^65,69–71^. Importantly, there is no cryo-EM information on MT-bound KIF1A dimers, the functional oligomeric form required for unidirectional processive motility^59^, especially with no structural data on MT-bound KIF1A in its two-heads-bound configuration. To fully understand the structural basis of KIF1A motility, high-resolution structures of KIF1A-MT complexes are needed. Such high-resolution data are essential to accurately trace the polypeptide chains, determine the position of side chains and differentiate between possible coexisting conformations. Furthermore, previous structural studies have largely left the functionally important K-loop unresolved, underscoring the necessity of high-resolution data not only for a comprehensive understanding of KIF1A’s motility and force-generation mechanisms but also for advancing structure-guided drug discovery efforts, which hinge on detailed insights into this key structural element.

To address these challenges, we determined the high-resolution cryo-EM structures of dimeric KIF1A bound to MTs at 2.7-3.4 Å overall resolution in various conformational states and nucleotide conditions: with the non-hydrolysable ATP analogue, AMP-PNP (adenylyl-imidodiphosphate), with ADP, and without nucleotides (Apo-state). These structures represent different stages of the KIF1A-MT ATPase cycle. Our results highlight the full structure of the K-loop and its interaction with the C-terminal tails of both α- and β-tubulin, providing functional evidence for the K-loop’s pivotal role in KIF1A superprocessivity. Moreover, we show structurally and functionally that the coordination between KIF1A’s two motor domains contributes to the processive motion of KIF1A. Additionally, we have determined the near-atomic resolution structure of the P305L KAND mutant bound to MTs, providing structural insights into how this mutation impairs KIF1A functionality.

In summary, our work provides critical insights into KIF1A’s mechanism and demonstrates the possibility of obtaining high-resolution structures of MT-bound KAND mutants, which could inform the development of targeted therapies based on these mutant structures.

## Results

### High-resolution structures of the KIF1A-MT complexes

To elucidate the structural basis of KIF1A’s unique motile properties, we obtained high resolution cryo-EM structures of dimeric KIF1A (Supplementary Fig. 1) bound to microtubules (MTs) in various nucleotide states, representing different stages of KIF1A’s ATPase cycle. We adapted a local classification and refinement strategy to the cryo-EM data^72^, which revealed distinct conformations of the two motor domains when associated with MTs (Supplementary Figs. 2 and 3). The cryo-EM maps achieved an overall resolution of 2.7-3.4 Å (Supplementary Fig. 4).

In the presence of AMP-PNP (abbreviated as ANP in our maps and model names), we observed classes where KIF1A’s two motor domains are connected by its coiled-coil dimerization domain and engaged with the core of two adjacent tubulin heterodimers along a MT protofilament (Fig. 1a). The leading head, positioned closer to the MT plus-end, features a backward-oriented neck-linker and an open nucleotide-binding pocket, whereas the trailing head exhibits a docked, forward-oriented neck-linker and a closed nucleotide-binding pocket. Both heads contain AMP-PNP bound in their nucleotide-binding sites (Figs. 1a and 2a). Other classes show single MT-bound heads with a trailing-like head and a missing leading head or with the trailing head of another dimer occupying the leading position (Supplementary Figs. 2 and 3). Our classification further revealed that the trailing head, whether in a two-heads-bound configuration or as a single-bound head with a docked neck-linker, adopts three different conformations (Supplementary. Fig. 3). Two classes, MT-KIF1A-ANP-T_2_L_1_ and MT-KIF1A-ANP-T_3_L_1_, shared strong similarities and were averaged to create the map depicted in Fig. 1a (MT-KIF1A-ANP-T_23_L_1_), from which we constructed the atomic model representing the average two-heads-bound configuration of KIF1A-ANP (Fig. 2a).

**Fig 1.**
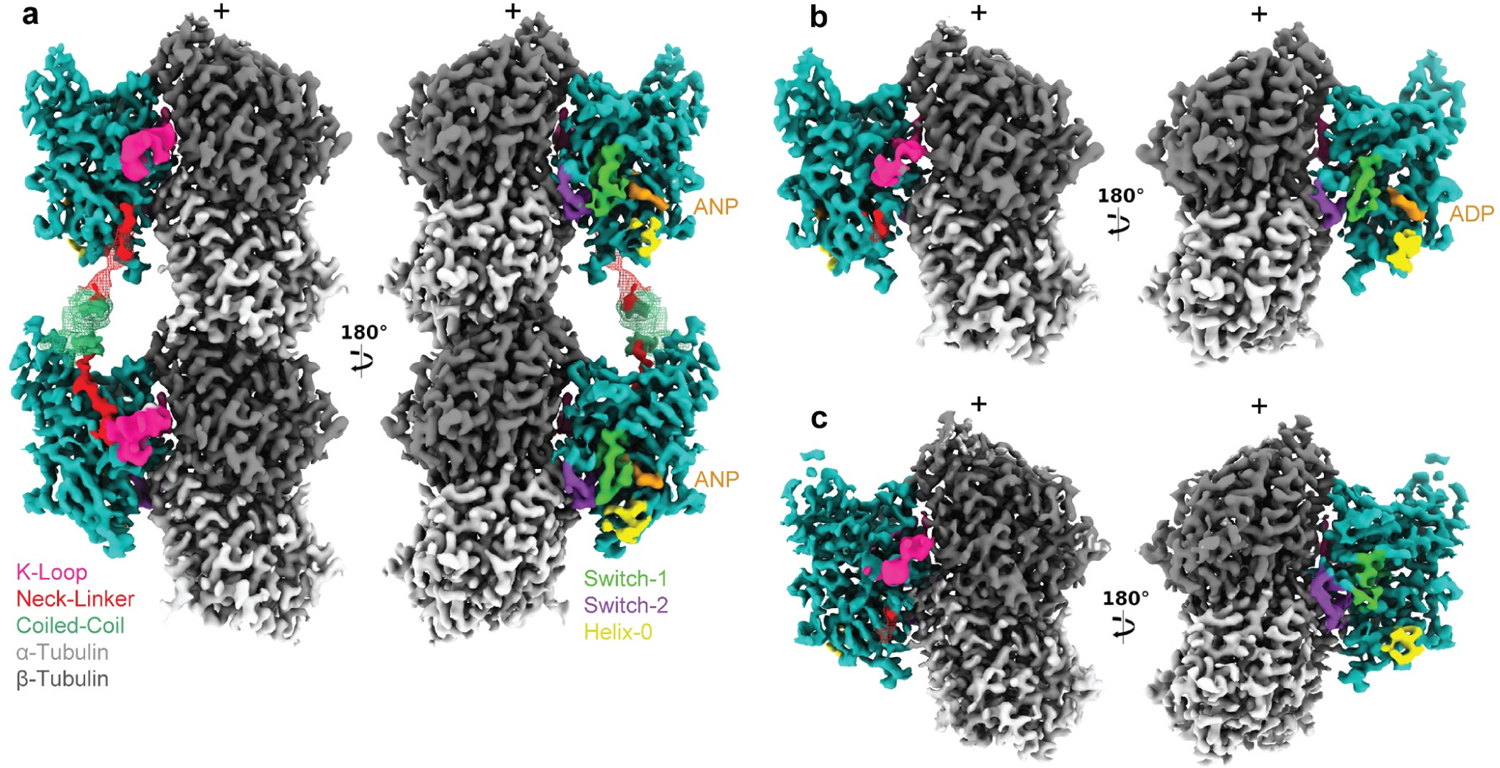
Cryo-EM maps of microtubule-bound KIF1A in different nucleotide states. Each panel shows two views of the isosurfaces of microtubule-bound KIF1A 3D maps, rotated 180° relative to each other. The surface colors, emphasizing different structural elements, as indicated in the figure labels. Map densities around the K-loop have been low-passed filtered and displayed at a lower contour level than the rest of the map for enhanced visualization of this mobile part. a: Microtubule-bound KIF1A with the ATP analogue AMP-PNP (abbreviated as ANP). The map corresponds to the average two-heads-bound configuration (MT-KIF1A-ANP-T_23_L_1_). The coiled-coil density and parts of the neck-linker from the leading head have been low-pass filtered and are displayed as a mesh at a lower contour level than that of the main map. Densities from the neck-linkers and coiled-coils connecting the two heads are visible. ANP densities are present in both the leading and trailing heads. Note: One-head-bound configurations were also found in the ANP datasets (Supplementary Fig. 2 & 3, Supplementary Table2). b-c: Isosurface representations of KIF1A in the ADP and APO states, (datasets MT-KIF1A-ADP and MT-KIF1A-APO).

**Fig. 2.**
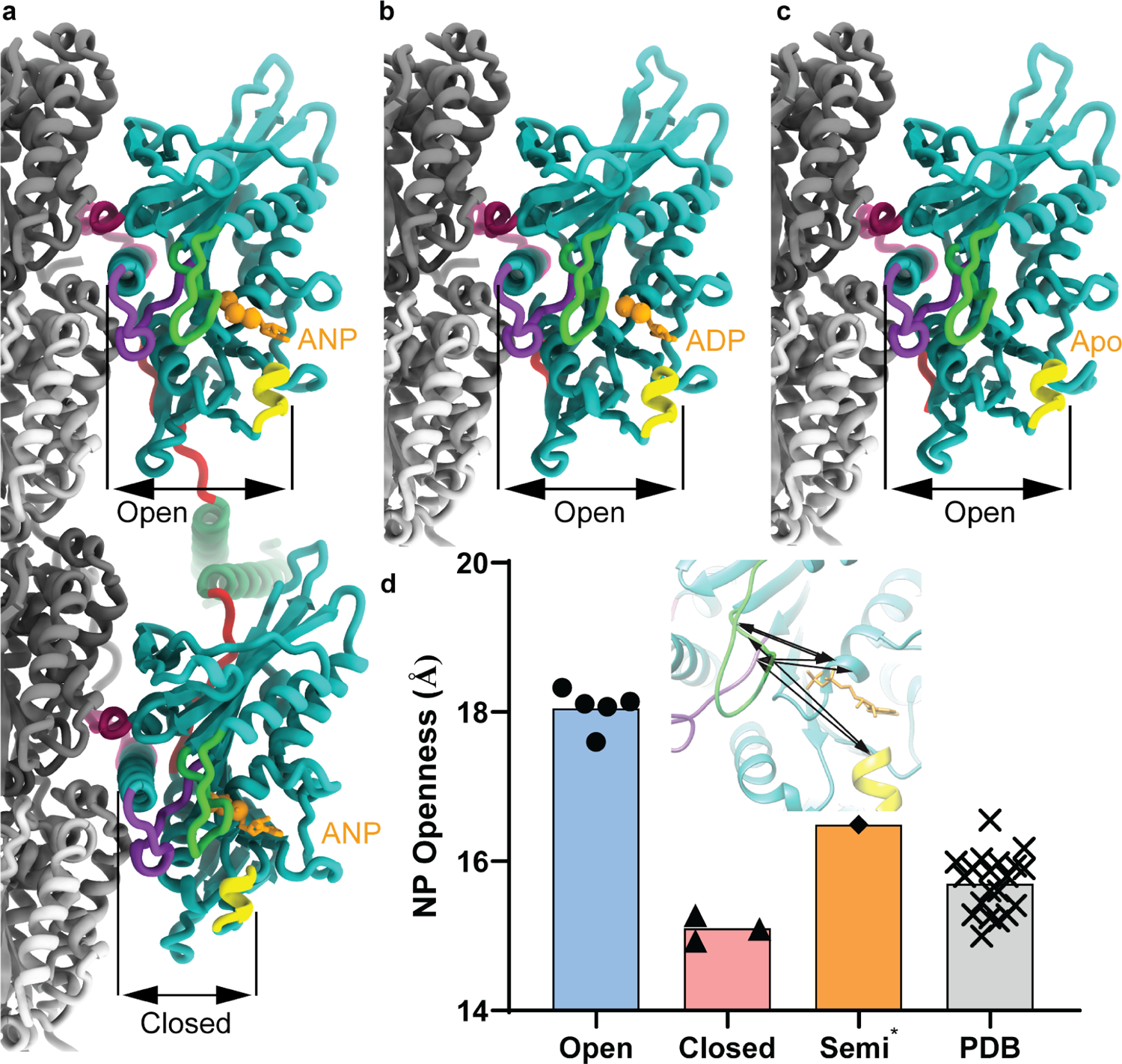
KIF1A atomic models. **a:** ANP two-heads-bound configuration (MT-KIF1A-ANP-T_23_L_1_). **b:** ADP one-head-bound configuration (MT-KIF1A-ADP). **c:** Apo one-head-bound configuration (MT-KIF1A-APO). **d:** Scatter plot illustrating the average of six distances between Cα carbons of selected residues (R216 and A250 to P14, to S104 and to Y105) across the KIF1A nucleotide-binding pocket (see inset). Circle symbols in the ‘Open’ column represent KIF1A motor domain structures in the ADP and Apo states, and the leading head in the two-heads-bound configuration of the ANP state. Triangle symbols in the ‘Closed’ column correspond to the structures of the trailing head in the two-heads-bound configuration of the ANP state. The diamond symbol in the ‘Semi^*’^ column represents a class structure of the ANP state in a single-head-bound configuration (MT-KIF1A-ANP-T_1_L_02*_). The cross symbols in the ‘PDB’ column represent KIF1A or KIF1A-MT models deposited in the PDB database, with accession codes: 1I5S, 1I6I, 1IA0, 1VFV, 1VFW, 1VFX, 1VFZ, 2HXF, 2HXH, 2ZFI, 2ZFJ, 2ZFK, 2ZFL, 2ZFM, 4UXO, 4UXP, 4UXR, 4UXS, 7EO9 and 7EOB. Bar height represents the mean openness value of the points in each column. NP: Nucleotide pocket.

**Fig. 3.**
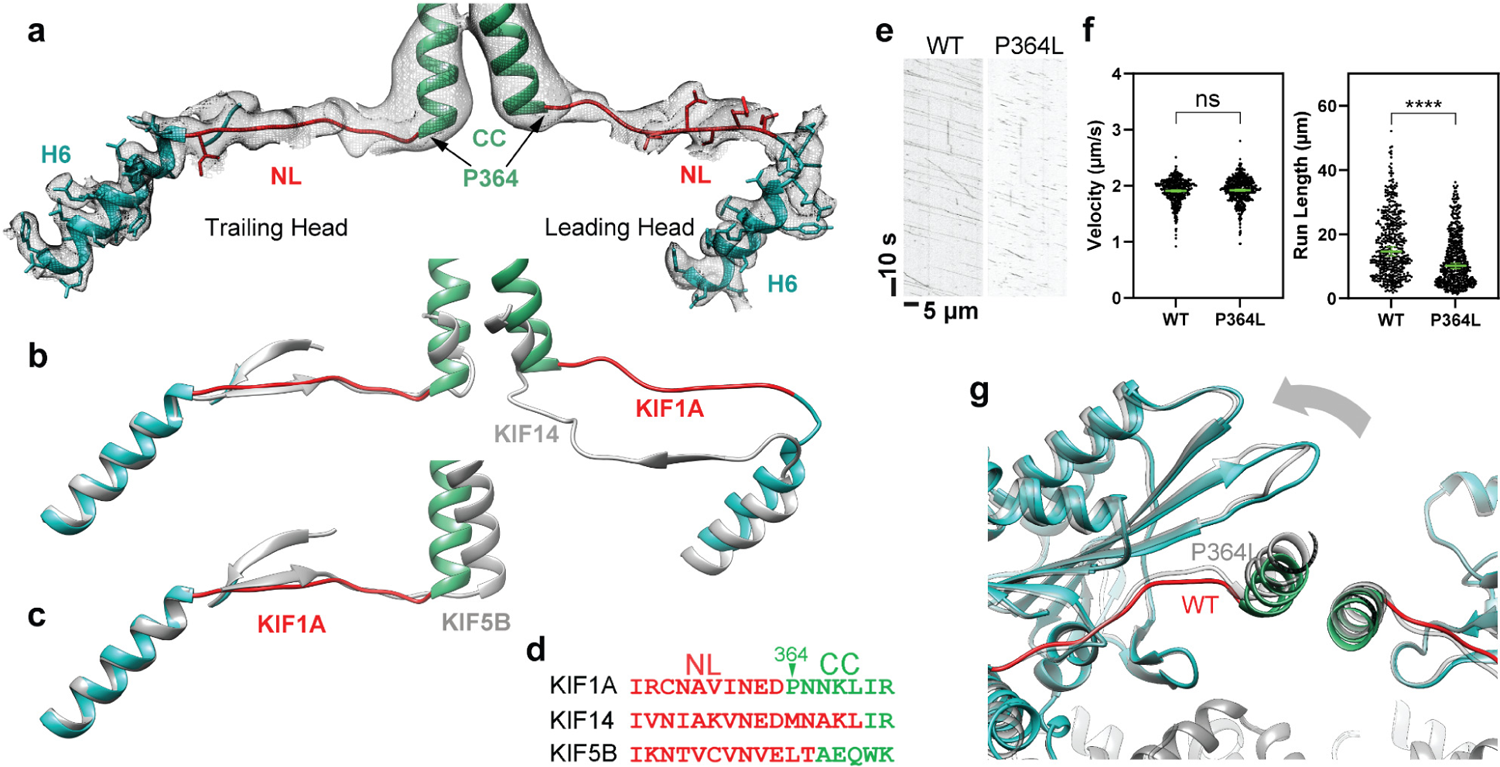
Neck-linker structure. **a:** H6, neck-linker (NL) and coiled-coil (CC) region of the KIF1A ANP two-heads-bound structure (MT-KIF1A-ANP-T_23_L_1_). The structure is presented as a ribbon representation with side chain atoms as sticks and color coded as in Fig. 1. The cryo-EM map is shown as an isosurface semitransparent mesh. A composite low pass-filtered version of the map highlights the NL and CC areas, with model side chains omitted. **b:** Comparison of the two-heads-bound configuration structures of KIF1A and KIF14 (grey color, PDB accession code: 6WWL). **c:** Comparison of KIF1A’s trailing head in the two-heads-bound configuration with KIF5B’s docked NL and CC helix crystal structure (shown in grey; PDB accession code: 1MKJ). **d:** Sequence alignment of NL and CC for KIF1A, KIF14 and KIF5B. Residues in the NL or the CC helix, as identified in structures (a) to (c), are colored red and green, respectively. **e:** Kymograph examples showing WT and P364L variant. **f:** Velocities and run-lengths of WT and P364L variant. Green bars represent mean values for velocity or median values for run length with their respective 95% confidence interval (CI). Velocities: WT: 1.91 [1.89, 1.93] µm/s, n=459; P364L: 1.92 [1.90, 1.94] µm/s n=514. Run lengths: WT: 14.6 [13.3, 16.1] µm, n=459; P364L: 10.2 [9.4, 11.0] µm, n=514. A minimum of three experiments were performed for each construct. Statistical analysis was conducted using an unpaired t-test with Welch’s correction (P=0.47) for velocities and Kolmogorov-Smirnov test for run lengths (****: P<0.0001). **g:** Superimposed ANP two-head-bound structure models of WT (MT-KIF1A-ANP-T_23_L_1_) and P364L (MT-KIF1A^P364L^-ANP-TL_1_), aligned by their tubulin components. The WT structure is colored blue and red, while the P364L variant is represented in gray.

The most distinct conformation, termed T1, was exclusively observed in one-head-bound classes and is characterized by having a docked neck-linker but a nucleotide-binding pocket that is semi-closed (Fig. 2d, Supplementary Figs. 3 and 5). This intermediate closure suggests that binding of the partner motor domain in the leading position may stabilize the fully closed conformation of the trailing head. This supports the notion of enhanced inter-head coordination, where the trailing head assumes a catalytically competent, closed conformation once its partner is bound to the MT in the leading and open conformation.

In conditions with ADP or without nucleotides (Apo state), only classes with single MT-bound heads were detected, lacking a visible connection to their partner head domains (Fig. 1b,c). This configuration represents a state where one head is MT-bound while the tethered partner head is unbound and mobile. Extra densities near the MT-bound motor in both Apo and ADP states hint that the unbound head partially explores the space behind the bound motor domain (Supplementary Fig. 6), as also suggested by the backward orientation of the linker’s initial segment in these structures (Supplementary Fig. 6). The structure of the MT-bound motor domain was similar in both conditions except for the presence or absence of ADP in the nucleotide-binding pocket (Figs. 1b,c and 2b,c). Notably, in the KIF1A ADP dataset, about 36% of the signal near the tubulin was unassigned, compared to a maximum of 5% in the other datasets (Supplementary Table 2). This significant portion of weak signal could represent weakly-bound mobile motors, consistent with the expected lower MT affinity in the ADP state.

The open and closed conformations of the nucleotide-binding pocket in MT-bound kinesins^72,73^ can be characterized by the distance between key residues involved in nucleotide binding (Fig. 2d). The largest and shortest distances values define respectively the open and closed conformations and intermediate distance values define the semi-closed (or semi-open) conformation. This conformation is observed in the crystal structures of MT-unbound free kinesins with ADP in the nucleotide-binding pocket^74^, including for KIF1A (Fig. 2d)^65^. The open conformation we observed in high-resolution KIF1A-MT complexes, both in ADP and Apo states and in the leading head with bound AMP-PNP, represents a novel conformation for the KIF1A motor domain. Previous X-ray crystallography-based models or models of KIF1A-MT complexes derived from lower resolution (>6 Å) cryo-EM data, show KIF1A in either the closed or semi-closed conformation (Fig. 2d)^65–68,70,71^. The discernible differences between our findings and earlier cryo-EM results likely stem from the higher resolution of our cryo-EM data, which enables us to distinguish between open and semi-closed conformation that would otherwise appear similar at lower resolutions. Our comparison between the crystal structures and the high-resolution KIF1A-MT complexes shows that MT binding directly induces the nucleotide-binding pocket’s transition from a semi-closed to an open conformation, providing a structural explanation for the enhanced ADP release associated with kinesin-MT binding^75–77^.

### Conformation of the KIF1A neck-linker

In the one-head-bound configuration of KIF1A observed in the presence of ADP and in the Apo state, the neck linker is undocked and extends backward towards the MT minus-end (Fig. 1 b, c, Supplementary Fig. 6e,f). Beyond the fourth neck-linker residue (N357), cryo-EM densities become notably weaker, indicating increased mobility in this region. In contrast, in the two-heads-bound configuration (MT-KIF1A-ANP-T_23_L1), clear cryo-EM densities reveal the neck-linkers of both trailing and leading heads, extending from the motor domain to the connecting coiled-coil dimerization domain (Figs. 1a, 3a). In this configuration, the neck-linker of the trailing head is docked onto the motor domain, while the neck-linker of the leading head remains undocked and oriented backward. This is also observed in the two-heads-bound MT complexes of KIF14^72^, the only other high-resolution structure of a kinesin in such a configuration currently available. However, distinct differences are evident in the KIF1A structure: the backward-oriented neck-linker of the leading head deviates more from the motor domain, taking a more direct, linear path between the two motor domains (Fig. 3a,b). These differences are likely due to the unique neck-linker sequences of KIF1A compared to other kinesins. Notably, KIF1A possesses a conserved proline at position 364, typical in the kinesin-3 family but absent in KIF14 and other kinesins (Fig. 3d). Within the two-heads-bound structure of KIF1A, this proline marks the start of an α-helix that pairs with a partner α-helix to form the coiled-coil dimerization domain (Fig. 3a). This feature results in a neck-linker for KIF1A that is shorter by approximately two to four residues compared to the neck-linkers of KIF14^72^ or the kinesin-1 motor KIF5B^78^ (Fig. 3b,c). Considering that the length of the neck-linker influences kinesin processivity^79^, we hypothesize that the shorter neck-linkers of KIF1A in the two-heads-bound conformation facilitate tighter coordination between the catalytic cycles of the two motor domains, thereby enhancing processivity.

To test whether the presence of a proline residue at the end of the KIF1A neck-linker affects its structure and KIF1A processivity we created a KIF1A variant by replacing the conserved proline residue with leucine (P364L). This substitution mirrors the residue found at the corresponding position in human KIF5B (L335) and in other kinesins (Fig. 3d). We then assessed the velocity and run-length of this variant and determined its two-heads-bound structure (Fig. 3e-g). Interestingly, the P364L mutation did not affect the motor’s velocity (Fig. 3e,f). However, it resulted in a modest but significant reduction in run length (from a median length of 14.6 µm to 10.2 µm, Fig. 3e,f), highlighting the crucial role of the proline residue in KIF1A’s superprocessivity.

The structural analysis of the P364L mutant revealed a neck-linker length similar to that of the wild-type (WT) (Fig. 3g). Nonetheless, replacing the proline likely increases conformational entropy in the substitution zone. Subtle variations in the neck-linker path and the orientation of the trailing motor domain are visible in the mutant (Fig. 3g). Such structural deviations hint at diminished interhead-tension in the P364L mutant, allowing a part of the trailing motor domain to shift slightly backward (Fig. 3g). Taken together, our functional and structural analyses suggest that the tight neck-linker connection between the two motor domains, as observed in the two-heads-bound configuration, plays a crucial role in KIF1A’s enhanced processivity.

### Structure and role of the KIF1A K-loop

The specific role of KIF1A’s K-loop in MT binding has been previously established^57,80–82^, yet its interaction mechanism with MTs has remained unclear. Previous X-ray crystallography and lower-resolution cryo-EM studies have not captured the conformation of this loop and how it interacts with MTs^65–68,70,71^. However, our high-resolution structures of KIF1A-MT complexes reveal the complete polypeptide path of the K-loop (Fig. 4a). Despite the challenges in identifying the exact side-chain positions due to lower resolution, we were able to delineate the K-loop’s overall trajectory. In each structure, the K-loop projects outward from the motor domain, situated opposite the nucleotide-binding site, nestled between KIF1A’s helix-4 and the β-tubulin C-terminal helix (helix-12).

**Fig. 4.**
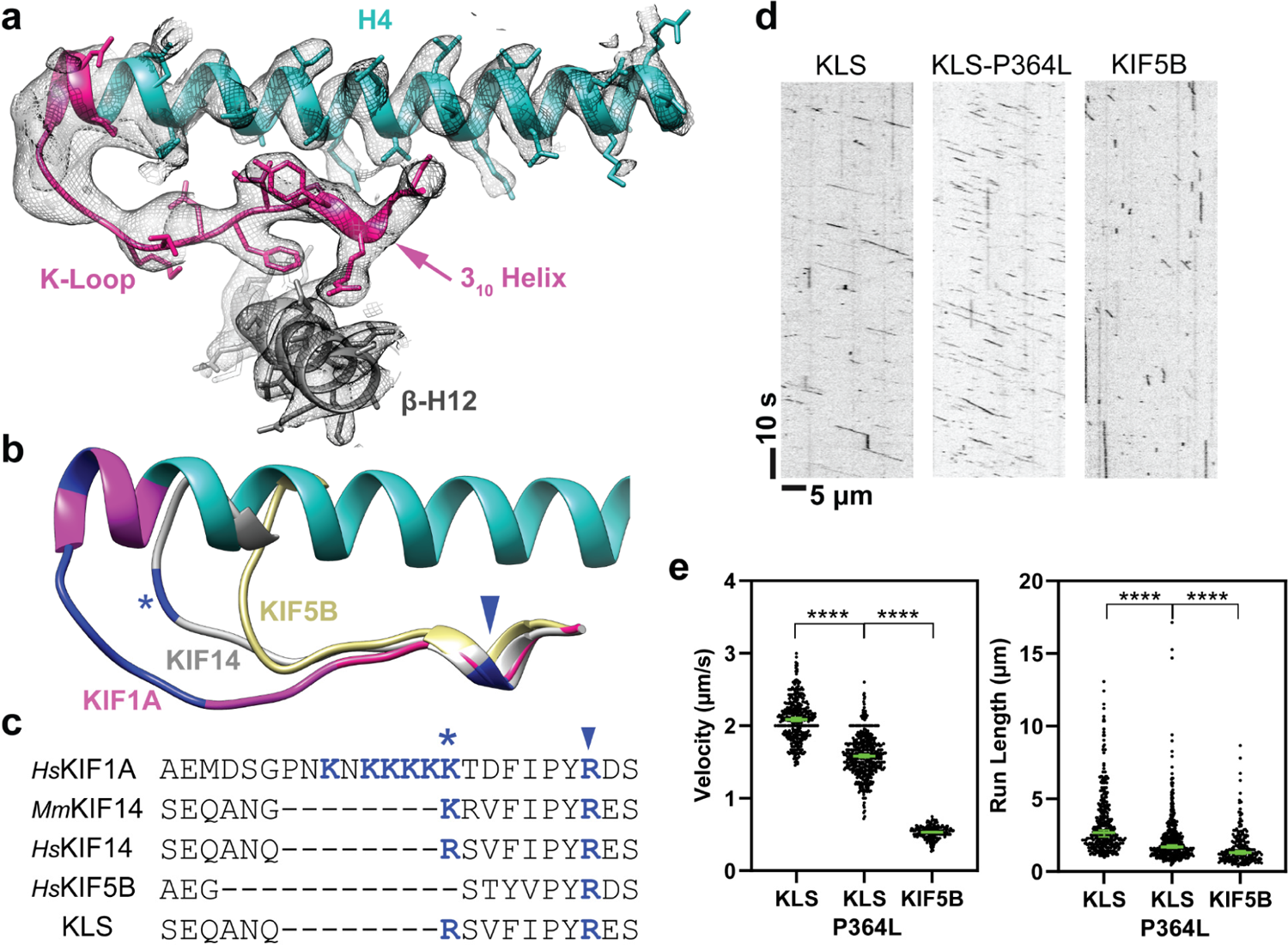
K-loop structure. **a:** Area of the K-loop of the KIF1A-ADP model. The model is shown as a ribbon representation with side chain atoms as stick. Parts colored as in Fig. 1. Cryo-EM density map is represented as a semi-transparent isosurface mesh, with a low pass-filtered version of the map emphasizing the K-loop tip area (left-most section). Model side chains are omitted in this area. **b:** Structural comparison of K-loop regions in *H*sKIF1A, *Mm*KIF14 (PDB accession code: 6WWM) and *Hs*KIF5B (PDB accession code: 1MKJ). Positively-charged residues (K or R) within the K-loop are highlighted in dark blue. A conserved positively-charged residue in kinesin-3’s loop12 is marked with a * symbol, and a highly conserved R residue in the kinesin superfamily is indicated by an arrowhead. **c:** Sequence alignments of loop-12 across different kinesins, and K-loop-swap mutant (KLS). Positively-charged residues are colored dark blue, with * and arrowhead symbols denoting the same residues marked in the structure (b). **d:** Kymograph examples of KLS, KLS-P364L and KIF5B. **e:** Velocities and run lengths of KLS, KLS-P364L and KIF5B. Green bars represent the mean values for velocity or median values for run length with their respective 95% CIs. Velocity: KLS: 2.09 [2.06, 2.11] µm/s n=398; KLS-P364L: 1.58 [1.56, 1.61] µm/s n=517; KIF5B: 0.53 [0.52, 0.54] µm/s n=232. Run-lengths: KLS: 2.7 [2.4, 2.9] µm n=398; KLS-P364L: 1.7 [1.6, 1.8] µm n=517; KIF5B: 1.3 [1.2, 1.5] µm n=232. A minimum of three experiments were performed for each construct. Statistical analysis was conducted using an unpaired t-test with Welch’s correction for velocities and Kolmogorov-Smirnov test for run lengths (****: P<0.0001).

The K-loop’s N-terminal lysines (K299 and K300) lie close to the β-tubulin’s terminal helix (Helix-12), suggesting potential electrostatic interactions with β-tubulin Asp-427. Interestingly, other kinesin-3s, like KIF14 (Fig. 4b,c), also possess positively charged residues in this area, albeit with a less extensive loop-12^72^. This suggests that the electrostatic interactions between this loop-12 region and the MT surface may play an important role in KIF1A motility.

To probe this idea, we engineered a chimeric construct where KIF1A’s K-loop was replaced with *Hs*KIF14’s loop-12 (K-loop-swap or KLS mutant, see methods). In near-physiological ionic strength buffer (BRB80, see methods), this construct exhibited a slightly higher velocity compared to WT KIF1A (Fig. 4d,e). However, the run-length was significantly shorter (reducing from a median of 14.6 µm to 2.7 µm), emphasizing the unique functional role of KIF1A’s K-loop in facilitating long run-lengths. This finding aligns with previous studies linking run length to the number of positively charged residues in the K-loop^82^.

Although the K-loop-swap mutant displayed a considerably shorter run-length, its median value still exceeded that of KIF5B (1.3 µm). Given that both proline 364 in the KIF1A neck-linker and the K-loop are distinct structural elements, we investigated whether combining these alterations would further reduce KIF1A’s processivity to match that of KIF5B. Introducing the double mutant (P364L-Kswap) resulted in a more substantial decrease in run length (1.7 µm) than either the P364L (10.2 µm) or K-loop-swap (2.7 µm) mutants individually. However, the run length of this double mutant still marginally surpassed that of KIF5B, suggesting additional factors in the motor domain contribute to KIF1A’s superprocessivity. This is in line with previous research showing that mutations of other KIF1A-specific residues at the MT interface lead to reduced processivity^58^. In terms of translocation velocity, both the P364L and the K-loop-swap mutants either maintained or slightly increased speeds relative to WT KIF1A. Conversely, the double mutation led to a 24% decrease in velocity compared to WT, highlighting a synergistic effect between P364 in the neck-linker and the K-loop in regulating KIF1A motility.

### Interactions of the K-loop with tubulin tails

Our cryo-EM maps detected densities close to the C-terminal tails of both α-tubulin and β-tubulin that extend into the K-loop region (Fig. 5). These densities appear less distinct and weaker compared to other areas in the cryo-EM maps, indicating increased mobility. The observed interactions of the tubulin C-terminal tails with the KIF1A K-loop provide a structural basis for the observation that removal of the tubulin C-terminal tails or the KIF1A K-loop significantly impact MT-binding and motility^57,79–82^.

**Fig. 5.**
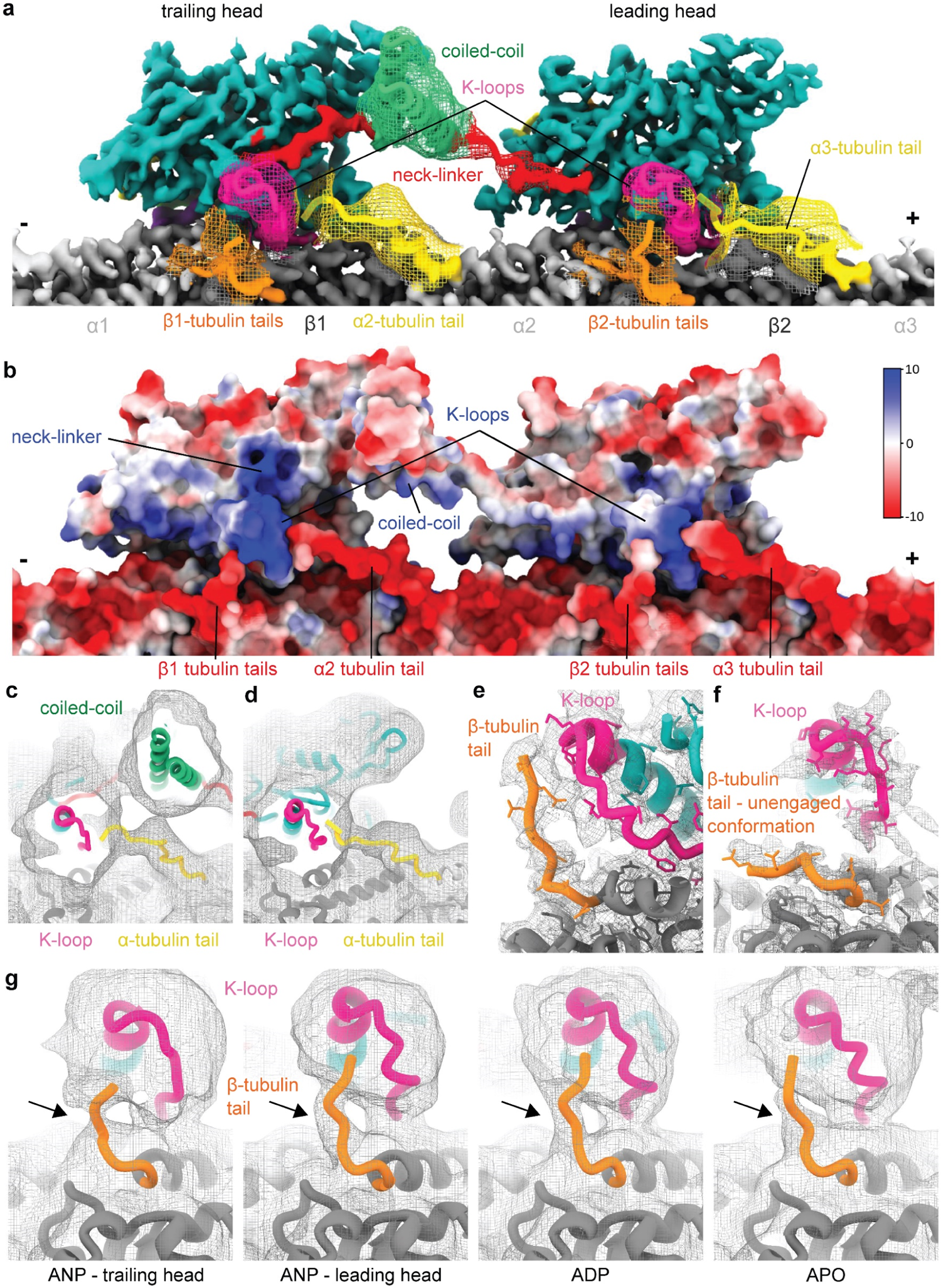
Interaction of the K-loop with the C-terminal tubulin tails. **a:** Composite cryo-EM map of KIF1A microtubule two-heads-bound configuration (MT-KIF1A-ANP-T_23_L_1_) interacting with three tubulin dimers. The microtubule is displayed horizontally throughout this figure with the plus end to the right. The locally filtered map is displayed as a solid color. The portions with weaker signals were low pass filtered and displayed as meshes with the underlying model displayed as a ribbon representation. The coiled-coil and β-tubulin tails interacting with the kinesin were low-passed filtered to 6 Å, the α-tubulin tail densities were low-passed filtered to 8 Å and displayed at a low-density threshold. Microtubule polarity is indicated, and the tubulin subunits are numbered from 1 to 3. The 6 Å and 8Å low-passed filtered maps used throughout this figure were deposited as additional maps in the corresponding EMDB entries. **b:** Representation of the Coulombic electrostatic surface potential of the KIF1A two-heads-bound configuration model viewed like in (**a**). Color scale in kcalꞏmol^-1^ꞏ*e*^-1^. **c-d:** 8 Å low-passed filtered cryo-EM map of KIF1A two-heads-bound configuration (MT-KIF1A-ANP-T_23_L_1_), showing densities of α-tubulin C-terminal tails reaching the K-loops of the trailing (**c**) or leading (**d**) head. The α-tubulin C-terminal tail densities were only detected in the ANP state. Note from panel (**b**) that the α-tubulin tail interacting with the trailing head is located within a pocket of positively charged residues due to the charges of K-loop, the nearby coiled-coil and the docked neck-linker. Given the shape of the map in this area it is likely that the α-tubulin tail interacts with all these KIF1A regions. **e:** Cryo-EM map of the leading head of the P364L mutant (MT-KIF1A^P364L^-ANP-TL_1_) showing a mostly resolved β-tubulin tail conformation interacting with the K-loop. **f:** Cryo-EM map of the trailing head of KIF1A two-heads-bound configuration (MT-KIF1A-ANP-T_23_L_1_) showing a portion of a β-tubulin tail conformation lying along the microtubule surface unlike the other bent conformation that interact with the K-loop as shown in (**e**) and (**g**). **g:** 6 Å low-passed filtered map of KIF1A near the K-loop in distinct nucleotide states and motor-domain conformations. The density associated with the β-tubulin C-terminal tail is pointed with an arrow and the threshold was adjusted for each map. Note that in the ANP state the β-tubulin tail density is better resolved in the leading head than in the trailing head. Comparisons of the β-tubulin tails of the leading and trailing head displayed at the same density threshold for the KIF1A WT and P364L mutant in the ANP state are given in Supplementary Fig. 7a-b.

In the case of the C-terminal tail of β-tubulin, we identified at least two concurrent positions: one aligned with the MT surface and another extending towards the K-loop (Fig. 5a,e,f). The coexisting positions may be due to high mobility or may also be a consequence of the presence of different post translational modification or tubulin isotypes in the MT. KIF1A interactions with the β-tubulin C-terminal tail were detected in all nucleotide states (Fig. 5g).

We also observed variations in the densities of the C-terminal tubulin tails, influenced by both the nucleotide state and the positioning of the motor domains (Fig. 5). Densities associated with the C-terminal tail of α-tubulin were only evident in the AMP-PNP state (Fig. 5a-d). Two factors may account for the difference between nucleotide states. In the two-heads-bound configuration the α-tubulin tail may help or be needed to secure MT binding of the motor-domain to the leading position. In the case of the trailing head the motor domain with its docked neck-linker, the K-loop, and a segment of the coiled-coil form a pocket of positively charged residues (Fig. 5b, c). This pocket likely attracts the α-tubulin C-tail. Moreover, the connection between the K-loop and the C-terminal tail of β-tubulin was less pronounced in the trailing motor domain compared to its leading counterpart (Fig. 5g, Supplementary Fig. 7a-b). These nucleotide-dependent variations in the interaction between the two tubulin C-terminal tails and the KIF1A motor domain provides a plausible explanation for the previously reported bias toward the MT plus-end of a monomeric KIF1A construct during one-dimensional diffusion along the MT lattice^80,83^ (Supplementary Fig. 8).

In summary, our results highlight the evolutionary adaptations of KIF1A’s K-loop and neck-linker. Their combined properties significantly enhance MT affinity and inter-head coordination, contributing to KIF1A’s exceptional processivity.

### High-resolution structures of KIF1A-MT complexes with pathogenic P305L mutation

To advance structure-based drug development, we determined high-resolution structures of KIF1A harboring the KAND-associated P305L mutation in complex with MTs, using the same nucleotide conditions as those for MT-bound WT KIF1A (Fig. 6a-c). Achieving sufficient MT decoration with the P305L mutant for cryo-EM imaging required a higher kinesin-to-MT ratio and a buffer with lower ionic strength compared to conditions used for the WT protein (Supplementary Table 1). Despite these adjustments, the decoration level for the mutant was lower than that of WT KIF1A (Supplementary Table 2), consistent with our previous findings of the mutant’s diminished MT-binding affinity^84^.

**Fig 6.**
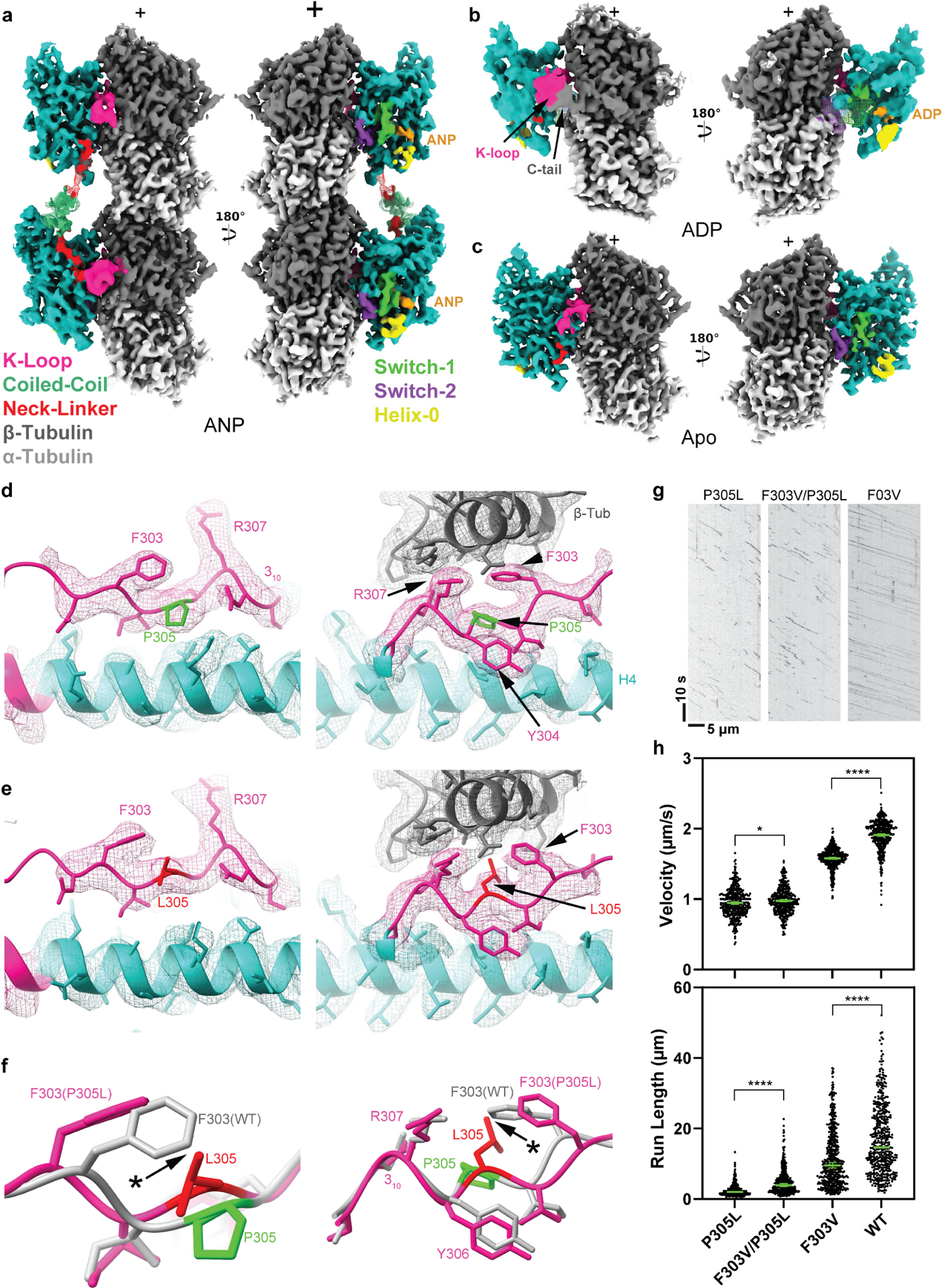
Cryo-EM maps of microtubule-bound KIF1A^P305L^ in different nucleotide states. Each panel presents two views of the isosurfaces of microtubule-bound KIF1A^P305L^ 3D maps, rotated 180° relative to each other. The surface colors, emphasizing different structural elements, as indicated in the figure labels (and as in Fig. 1). The K-loop’s signal has been low-passed filtered and displayed at a lower contour level than the rest of the map for enhanced visualization. **a:** MT-bound KIF1A^P305L^ with ANP. The map represents the two-heads-bound configuration MT-KIF1A^P305L^-ANP-TL_1_. The coiled-coil density and part of the neck-linker from the leading head were low-pass filtered and displayed as a mesh and at much lower contour level than the main map. Densities from the neck-linkers and coiled-coils connecting the two heads are visible. The P305L mutant in the ANP state exhibits both two-heads-bound (displayed here) and single-head-bound configurations (Supplementary Fig. 9, Supplementary Table 2), In the two heads-bound configuration the trailing head is in a closed conformation with a docked neck-linker and the leading head in an open conformation with the neck-linker oriented backwards. ANP densities are present in both the leading and trailing heads. **b:** Isosurface representation of KIF1A^P305L^ in the ADP state (dataset MT-KIF1A^P305L^-ADP). This lower resolution map displays the KiF1A switches area as a low-pass filtered mesh. The K-loop area and C-terminal tail of β-tubulin, with connected densities, are low-passed filtered and displayed as a solid surface. **c:** Isosurface representations of KIF1A^P305L^ in the Apo state (dataset MT-KIF1A^P305L^-APO), illustrated as in panel (**a**). Both motors KIF1A^P305L^ in the ADP and APO datasets are in the open conformation like KIF1A WT (Fig. 1-2). **d:** Isosurface representations of the cryo-EM map of WT KIF1A in the ANP state (MT-KIF1A-ANP-T_23_L_1_) in the neighborhood of residue P305. The surface is represented as a mesh and colored based on the underlying fitted molecular model (same color convention as in Fig. 1). The model is shown as ribbon representation with displayed side chains. The left and right panels represent two different views of the same area, the left one looking from β-tubulin, on an axis orthogonal to the microtubule axis, and the right one along the microtubule axis, from the (+) end towards the (-) end. All the side chains are resolved in this area, including P305, colored in green. **e:** Isosurface representations of the cryo-EM map of KIF1A^P305L^ mutant in the ANP state (MT-KIF1A^P305L^-ANP-TL_012*_), same view as in (**d**). The side chains are also well resolved in the KIF1A^P305L^, including L305, colored in red. L305 is positioned differently from P305, with L305 pointing more towards the microtubule interface. **f:** Overlay of the models displayed in (**d**) and (**e**) with the KIF1A WT model displayed in gray, the KIF1A^P305L^ mutant model in pink and the residue 305 colored as in panels (**d-e**). This overlay emphasizes the differences between KIF1A and KIF1A^P305L^ in the neighborhood of residue 305. In addition to a local backbone change at position 305 due to the absence of the proline in L305, the side chain of L305 would clash with the side chain of F303 in the WT configuration (pointed with the symbol * and the arrow). This causes a reorientation of F303 in the P305L mutant relative to WT. The changes up the polypeptide chain after 305 are more modest with the path of the 3_10_-helix being very similar, including the position of the highly conserved R307. **g:** Kymograph examples of P305L, F303V/P305L and F303V. **h:** Distribution of velocities and run lengths. The green bars represent the mean for velocity or median values for run length with their respective 95% confidence interval. Velocity: P305L: 0.95 [0.93, 0.96] µm/s, n=438; F303V/P305L: 0.98 [0.96, 0.99] µm/s n=441; F303V: 1.58 [1.57, 1.59] µm/s n=557; WT: 1.91 [1.89, 1.93] µm/s, n=459. The statistics were performed using unpaired t-test with Welch’s correction (****: P<0.0001; *: P=0.016). At least 3 experiments were performed for each construct. Run lengths: P305L: 2.0 [1.9, 2.2] µm n=438; F303V/P305L: 3.9 [3.5, 4.4] µm n=441; F303V: 9.5 [8.8, 10.3] µm n=557; WT: 14.6 [13.3, 16.1] µm n=459. The statistics were performed using Kolmogorov-Smirnov test (****: P<0.0001).

The KIF1A-P305L mutant, like its WT counterpart, exhibits classes with both one- and two-heads-bound configurations in the presence of AMP-PNP: the leading head in an open conformation and the trailing head in a closed conformation with nucleotide bound to both heads (MT-KIF1A^P305L^-ANP-TL_1_, Fig 6a, Supplementary Fig. 9). In the ADP and Apo states, it shows a one-head-bound open conformation (Fig. 6b,c). Weak densities corresponding to α- and β-tubulin tails connecting to the motor domain were detected in all nucleotide states (Supplementary Figure 7 c-h), suggesting that the P305L mutation does not prevent interactions between the K-loop and the α and β-tubulin tails. Notably, in the ADP state, the P305L mutant displays a distinct map with motor domain densities appearing at much lower resolution compared to the MT densities (Fig. 6b). The lower resolution can be attributed to increased mobility, rather than poorer decoration of the mutant in the presence of ADP because the classification method separates the motor-decorated tubulins from the undecorated tubulins, and the resolution of the MT part of the map is comparable to the MT resolution of other maps presented (Table 1).

**Table 1.** Cryo-EM data collection, refinement and validation statistics (1/3)

|  | **MT-KIF1A-**<br>**ANP-T<sub>23</sub>L<sub>1</sub>**<br>(EMDB-42543)(PDB<br>8UTN) | **MT-KIF1A-**<br>**ANP-T<sub>2</sub>L<sub>1</sub>**<br>(EMDB-42544)<br>(PDB 8UTO) | **MT-KIF1A-**<br>**ANP-T<sub>3</sub>L<sub>1</sub>**<br>(EMDB-42545)<br>(PDB 8UTP) | **MT-KIF1A-**<br>**ANP-T<sub>1</sub>L<sub>02</sub>\***<br>(EMDB-42546)<br>(PDB 8UTQ) |
| --- | --- | --- | --- | --- |
| **Data collection and processing** |  |  |  |  |
| Magnification (actual) | 59242 | 59242 | 59242 | 59242 |
| Voltage (kV) | 300 | 300 | 300 | 300 |
| Electron exposure (e<sup>-</sup>/Å<sup>2</sup>) | 50.2 | 50.2 | 50.2 | 50.2 |
| Defocus range (µm) <sup>a</sup> | 0.8 ; 1.7 | 0.8 ; 1.7 | 0.8 ; 1.7 | 0.8 ; 1.7 |
| Pixel size (Å) | 0.844 | 0.844 | 0.844 | 0.844 |
| Symmetry imposed <sup>b</sup> | Helical | Helical | Helical | Helical |
| Rise (Å) | 5.60 | 5.60 | 5.60 | 5.60 |
| Twist (deg) | 168.09 | 168.09 | 168.09 | 168.09 |
| Particle images identified as 15R symmetry (no.) | 60199 | 60199 | 60199 | 60199 |
| Particle images in helical reconstruction (no.) | 60199 | 60199 | 60199 | 60199 |
| Single particles (no.) <sup>c</sup> | 902985 | 902985 | 902985 | 902985 |
| Single particles used (no.) <sup>d</sup> | 117091 | 99325 | 89711 | 116194 |
| Overall resolution (Å) | 3.1 | 3.2 | 3.2 | 3.1 |
| FSC threshold | 0.143 | 0.143 | 0.143 | 0.143 |
| Kinesin resolution (Å) | 3.2 | 3.3 | 3.3 | 3.3 |
| Tubulin resolution (Å) | 3.0 | 3.1 | 3.1 | 3.0 |
| **Refinement** |  |  |  |  |
| Model composition |  |  |  |  |
| Non-hydrogen atoms | 23787 | 23791 | 23791 | 13356 |
| Protein residues | 2981 | 2982 | 2982 | 1680 |
| Ligands | 13 | 13 | 13 | 8 |
| R.m.s. deviations |  |  |  |  |
| Bond lengths (Å) | 0.0063 | 0.0052 | 0.0054 | 0.0053 |
| Bond angles (°) | 1.18 | 1.12 | 1.13 | 1.13 |
| Validation |  |  |  |  |
| MolProbity score | 1.73 | 1.40 | 1.52 | 1.51 |
| Clashscore | 4.92 | 3.38 | 4.14 | 3.99 |
| Poor rotamers (%) | 1.64 | 0.43 | 0.94 | 0.28 |
| Ramachandran plot |  |  |  |  |
| Favored (%) | 95.48 | 96.01 | 95.35 | 95.39 |
| Allowed (%) | 3.67 | 3.72 | 4.21 | 4.49 |
| Disallowed (%) | 0.84 | 0.27 | 0.44 | 0.12 |

<sup>a</sup> Range of the per-particle defocus estimates. This range encompasses the defocus values for 90% of the particles used, with 5% of the particle defocuses falling below and 5% exceeding this range.
<sup>b</sup> Symmetry is imposed during the initial symmetrical refinement preceding the local refinements.
<sup>c</sup> Total number of particles after symmetry expansion.
<sup>d</sup> Number of asymmetric units used after 3D classification, corresponding to a kinesin motor bound to a tubulin dimer.

**Table 1.**
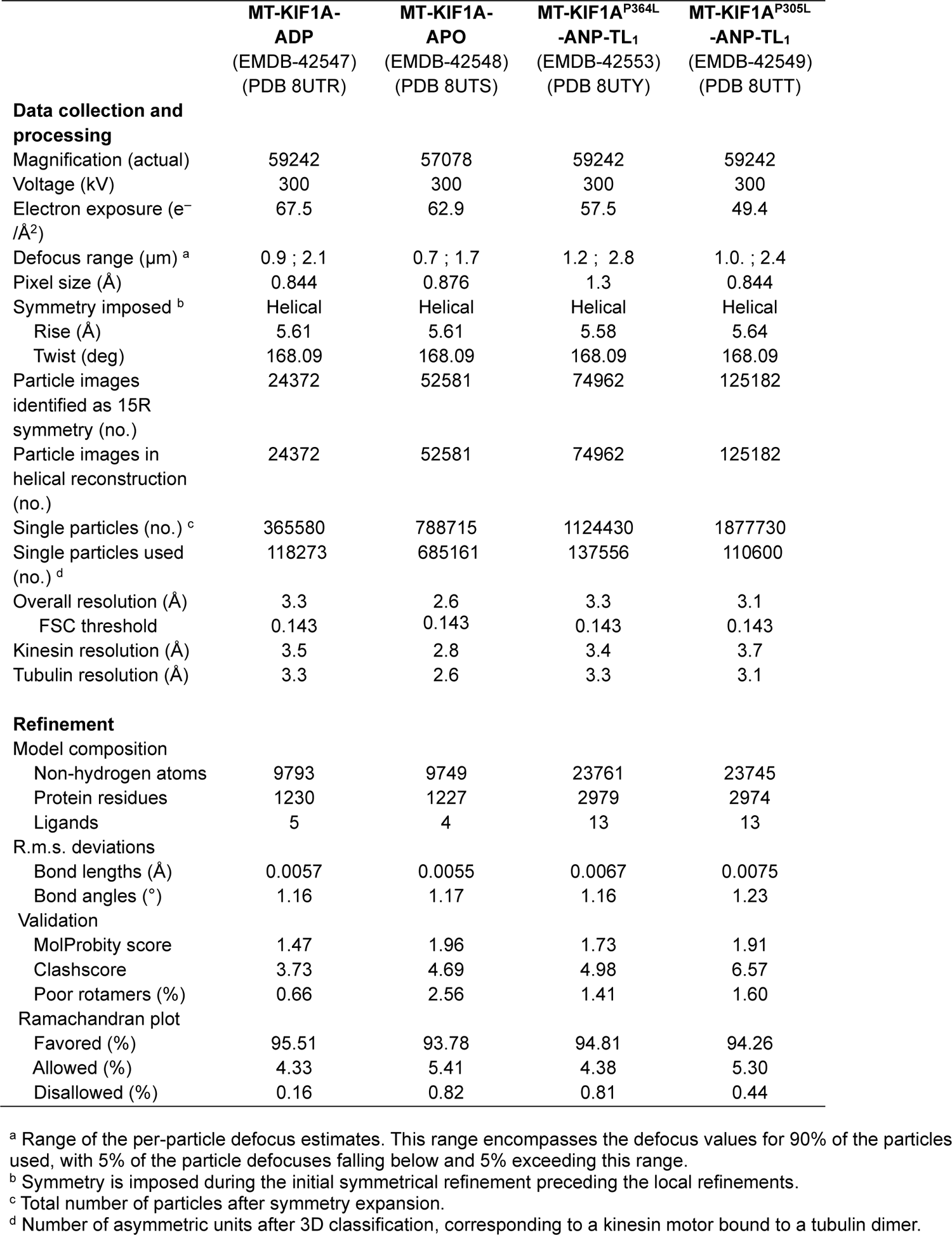
Cryo-EM data collection, refinement and validation statistics (2/3)

**Table x.** Cryo-EM data collection, refinement and validation statistics (3/3)

|  | **MT-KIF1A<sup>P305L</sup>**<br>**-ANP-TL<sub>012</sub><sup>\*</sup>**<br>**(EMDB-42550)**<br>**(PDB 8UTU)** | **MT-KIF1A<sup>P305L</sup>**<br>**-ADP**<br>**(EMDB-42551)**<br>**(PDB 8UTV)** | **MT-KIF1A<sup>P305L</sup>**<br>**-APO**<br>**(EMDB-42552)**<br>**(PDB 8UTW)** |
| --- | --- | --- | --- |
| **Data collection and processing** |  |  |  |
| Magnification (actual) | 59242 | 57078 | 59242 |
| Voltage (kV) | 300 | 300 | 300 |
| Electron exposure (e<sup>-</sup>/Å<sup>2</sup>) | 49.4 | 62.9 | 50.4 |
| Defocus range (μm) <sup>a</sup> | 1.0 ; 2.4 | 0.8 ; 2.0 | 1.2. ; 2.2 |
| Pixel size (Å) | 0.844 | 0.876 | 0.844 |
| Symmetry imposed <sup>b</sup> | Helical | Helical | Helical |
| Rise (Å) | 5.64 | 5.64 | 5.62 |
| Twist (deg) | 168.09 | 168.09 | 168.09 |
| Particle images identified as 15R symmetry (no.) | 125182 | 83861 | 79145 |
| Particle images in helical reconstruction (no.) | 125182 | 83861 | 79145 |
| Single particles (no.) <sup>c</sup> | 1877730 | 1257915 | 1187175 |
| Single particles used (no.) <sup>d</sup> | 254197 | 119590 | 620149 |
| Overall resolution (Å) | 3.0 | 3.0 | 3.4 |
| FSC threshold | 0.143 | 0.143 | 0.143 |
| Kinesin resolution (Å) | 3.4 | 3.8<sup>e</sup> | 4.1 |
| Tubulin resolution (Å) | 3.0 | 2.9 | 3.4 |
| **Refinement** |  |  |  |
| Model composition |  |  |  |
| Non-hydrogen atoms | 13617 | 13338 | 9787 |
| Protein residues | 1709 | 1679 | 1227 |
| Ligands | 8 | 7 | 4 |
| R.m.s. deviations |  |  |  |
| Bond lengths (Å) | 0.0064 | 0.0058 | 0.0142 |
| Bond angles (°) | 1.23 | 1.18 | 1.24 |
| Validation |  |  |  |
| MolProbity score | 1.67 | 1.53 | 2.02 |
| Clashscore | 5.26 | 4.76 | 7.73 |
| Poor rotamers (%) | 1.43 | 0.70 | 1.60 |
| Ramachandran plot |  |  |  |
| Favored (%) | 96.12 | 95.87 | 93.12 |
| Allowed (%) | 3.65 | 4.07 | 6.47 |
| Disallowed (%) | 0.24 | 0.06 | 0.41 |

<sup>a</sup> Range of the per-particle defocus estimates. This range encompasses the defocus values for 90% of the particles used, with 5% of the particle defocusses falling below and 5% exceeding this range.
<sup>b</sup> Symmetry is imposed during the initial symmetrical refinement preceding the local refinements.
<sup>c</sup> Total number of particles after symmetry expansion.
<sup>d</sup> Number of asymmetric units after 3D classification, corresponding to a kinesin motor bound to a tubulin dimer.
<sup>e</sup> Note that in this map kinesin density is weak and of low-resolution with no side-chains resolved unlike in the tubulin part of the maps. As a consequence, the kinesin resolution given is an overestimate (likely >>3.8 Å) dominated by the signal near the microtubule interface.

Despite the poorer resolution, densities between the tubulin tails and the KIF1A K-loop are observed (Fig. 6b), suggesting that this weakly MT-bound configuration is mediated mainly through electrostatic interactions between the K-loop and the tubulin tails. The low resolution of the kinesin part of that map prevented us from quantifying the openness of the nucleotide pocket, while the overall conformation appeared similar to that of WT KIF1A-ADP. We propose that the P305L mutation hinders the formation of a strongly MT-bound configuration, thereby increasing the fraction of weakly attached motor domains in the MT images. Supporting this hypothesis, the P305L-ADP dataset has the lowest level of kinesin decoration (∼9%) among all datasets (Supplementary Table 1).

To understand the structural impact of the P305L mutation on MT binding, we compared the mutant and WT KIF1A structures near the mutated P305L residue (Fig. 6d-f). P305 is a key component of the MT-binding interface, located within the highly conserved loop-12 motif, (PYRD/E), which forms a 3_10_-helix in various kinesins, including KIF1A^84^. Although previous assumptions suggested that the P305L mutation might disrupt this helix^84^, our findings indicate subtler conformational shifts (Fig.6d-f). The integrity of the 3_10_-helix remains intact, with most changes occurring towards the N-terminal side of the mutation. A notable divergence in the P305L mutant is the reorientation of F303 due to clash with the introduced leucine residue (Fig. 6f).

To assess whether the constrained position of F303 by L305 contributes to a reduced binding by impacting the MT-unbound KIF1A, we use the ColabFold^85^ implementation of AlpfaFold2 (AF2)^86^ to model WT KIF1A and the P305L mutant (Supplementary Fig. 10). The resulting 5 best AF2 models of WT KIF1A closely resembled the MT-bound structures near L305 (Supplementary Fig. 10a,c,e). However, in the P305L mutant models with the highest score, F303 is positioned in a way that would severely clash with the β-tubulin helix H12, rendering it incompatible with MT binding (Supplementary Fig. 10b,d,f). In some P305L models, part of the loop-12 forms a α-helix absent in the WT AF2 models and MT-bound structures. These results suggest that the P305L mutation induces conformations suboptimal for MT binding of F303 and other loop-12 areas in the bound and unbound states. Consequently, this mutation likely shifts the equilibrium towards the MT-unbound (or weakly-bound) states, as we observe here.

The structural differences observed between the P305L mutant and WT KIF1A suggested the possibility of restoring the functionality of the P305L mutant by introducing a residue at position 303 that would not clash with the leucine at position 305 of the mutant. To test this hypothesis, we analyzed the motility of a KIF1A F303V single mutant and a F303V/P305L double mutant (Fig. 6g-h). We selected valine because of its smaller size and hydrophobic character, and also because it occurs naturally at this position in some kinesin-3s, such as Unc-104. The F303V single mutant displayed reduced velocities and run lengths compared to WT KIF1A. Intriguingly, the double mutant exhibited an increased run length in comparison to the P305L mutant alone, indicating a partial rescue of processivity in the double mutant.

## Discussion

KIF1A, a key member of the kinesin-3 family, is recognized for its high processivity and fast MT plus-end-directed motility. These attributes are crucial for the long-distance transport of cargoes in neurons^3–5^. Despite its importance, the underlying mechanisms of KIF1A’s distinctive motility properties have remained largely undefined. Notably, most of the approximately 150 KAND mutations are within the motor domain^9^, yet high-resolution structures of MT-bound KIF1A were previously unavailable and intermediates in the mechanochemical cycle such as the two-heads-MT-bound configuration had never been observed. This lack of detailed structural information, particularly concerning intermediates like the two-heads-MT-bound configuration, has hindered a deeper understanding of KIF1A’s motility mechanism and limited the scope of structure-based drug discovery efforts for KAND. In this study, we have bridged this knowledge gap by determining high-resolution structures of dimeric, MT-bound KIF1A across various nucleotide states and by solving the first high-resolution structures of the KAND-associated P305L mutant. These structures have revealed previously unseen structural features essential for KIF1A’s motility mechanism, including a novel open conformation of the nucleotide-binding pocket, the MT-bound conformation of KIF1A’s K-loop, the dimeric structure of KIF1A in a two-heads-bound configuration, and the structural changes in the motor domain caused by the KAND P305L mutation.

In the ADP or Apo state, KIF1A is characterized by a single MT-bound motor domain, wherein the nucleotide-binding pocket adopts a markedly more open conformation than what has been documented in previous ‘free’, MT-unbound crystal structures of KIF1A-ADP^65,67^. This open conformation is reminiscent of the ones observed in high-resolution structures of tubulin- and MT-bound kinesin complexes from various subfamilies^72,73,87,88^. Thus, the open conformation identified in our KIF1A-MT complexes underscores the existence of a conserved mechanism throughout the kinesin superfamily to couple MT binding with product release^71^.

In previous structural studies, the K-loop of KIF1A has consistently remained unresolved^65,69–71^, leading to a question about whether its elusiveness was due to intrinsic disorder, mobility of the loop, limitations of cryo-EM resolution, or a combination thereof. Our high-resolution structures of KIF1A-MT complexes now reveal the complete polypeptide path of the K-loop, although with a resolution that is reduced relative to other regions, suggesting higher mobility in this area. These findings imply that earlier limitations in observing the K-loop may have been related to the resolution limits of cryo-EM at the time. However, the persistent absence of a fully resolved K-loop in high-resolution of X-ray crystallography structures suggests that the loop possesses inherent disorder in the MT-unbound KIF1A. Upon MT binding, we observe that the K-loop attains a degree of structural order, becoming discernible in the context of its association with MTs. This ordering upon MT binding echoes patterns seen in other kinesins, where interaction with MTs induces structuring in parts of the motor domain that form the MT-binding interface^70,72,89,90^.

Our study illuminates the structural positioning of the K-loop in KIF1A, revealing that while the bulk of the loop, including its key lysine residues, is distanced from the main body of the MT, it engages in dynamic electrostatic interactions with the C-terminal tails of both α- and β-tubulin. The K-loop effectively acts as an electrostatic bridge sandwiched between the two tubulin tails. This observation provides a structural basis for understanding the reduced run length we observed when we replaced KIF1A’s K-loop with the loop-12 of the kinesin-3 KIF14, which contains a single positively-charged residue near the MT interface. This finding aligns with recent studies, highlighting the crucial role of the K-loop and its positively-charged residues in facilitating KIF1A’s superprocessivity^81,82^.

Our structural and functional insights indicate that the K-loop’s impact on KIF1A’s superprocessivity primarily stems from dynamic electrostatic interactions between the K-loop and the MT. Such connections offer a structural foundation for a highly dynamic MT-bound configuration, where a single motor domain executes directionally biased one-dimensional diffusion along the MT lattice^83^. This phenomenon has been shown to rely on the K-loop and the C-terminal tubulin tails^80^. Moreover, we observe nucleotide-dependent interactions between the α- and β-tubulin C-terminal tails with the K-loop and possibly the coiled-coil domain. This provides a plausible mechanism for the reported plus-end-biased, one-dimensional diffusion of monomeric KIF1A^83^ (Supplementary Fig. 8).

Our structural findings can be integrated into an updated model for KIF1A’s processive motion along MTs, which aligns with our results and prior observations (Fig. 7). As in other models^75,91^, it depicts the two motor domains alternating between MT-bound and unbound configurations. The model incorporates that the activity of both motor domains is coordinated to ensure that their ATPase cycles remain ‘out-of-phase,’ which effectively reduces the likelihood of both heads assuming simultaneously the low-affinity ADP state and preventing detachment from the MT. The KIF1A-MT structures provide evidence for at least three points of coordination between motor domains in the KIF1A mechanochemical cycle (Fig. 7 legend). In addition, in scenarios where only one head is bound or both heads transiently enter the low-affinity ADP state, the electrostatic interactions between the C-terminal tubulin tails and KIF1A’s K-loop reduce the likelihood of complete detachment and may facilitate transition to a strongly MT-bound configuration, which are critical features for KIF1A’s superprocessive motion.

**Fig 7.**
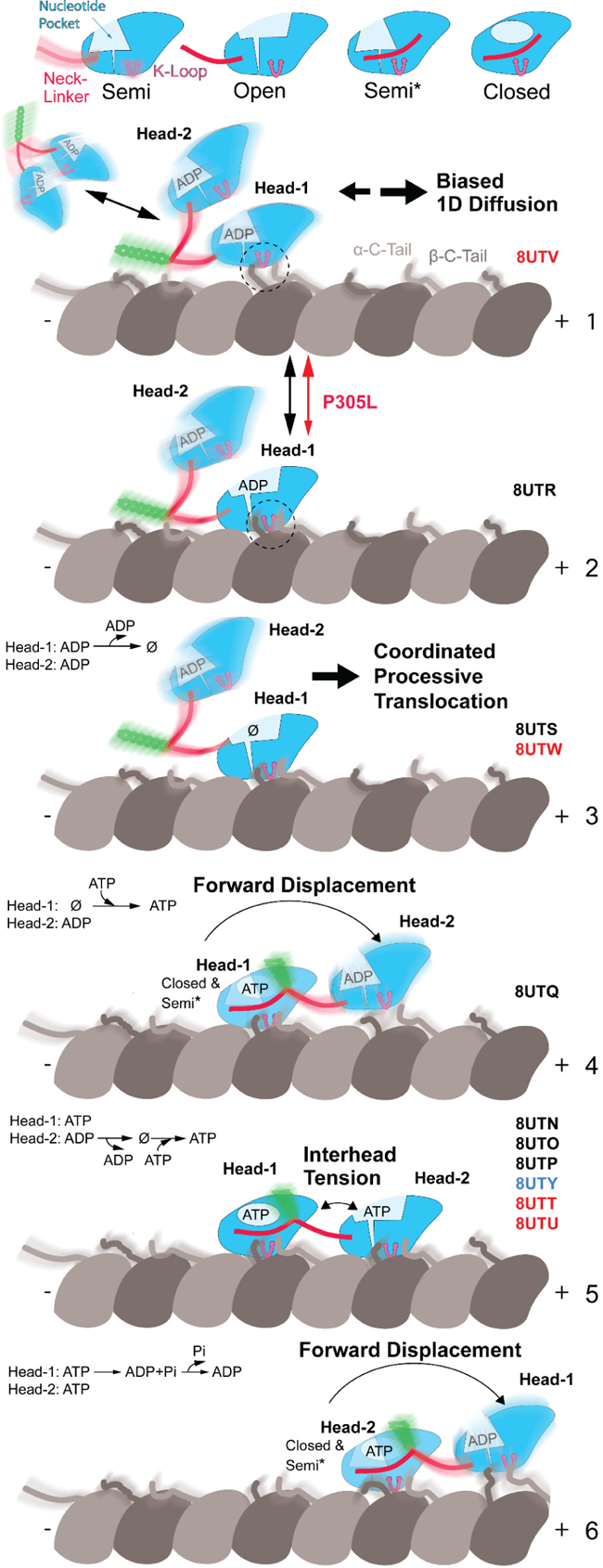
A model for KIF1A’s mechanochemical cycle. The model is based on the KIF1A-MT structures bound to different nucleotide analogues. The upper row displays the four primary conformations of the KIF1A motor domain, named based on the degree of openness of the nucleotide-binding pocket (Fig. 2d and Supplementary Fig. 5). The Semi (or semi-closed) conformation corresponds to the ADP-bound KIF1A motor domain crystal structures and the others corresponds to MT-bound structures determined in this work. The Open conformation corresponds to the MT-bound structures in the ADP and Apo states and the leading head of a two-heads-bound configuration in the ANP state (ATP bound state). The Closed conformation corresponds to two slightly different classes of the trailing head of the ATP-bound state (T2 and T3). The Semi* conformation has a semi-closed nucleotide pocket and a docked neck-linker. This conformation corresponds to a class (T1) observed in the ATP-bound state in a single-bound-head configuration. Two types of KIF1A-MT interactions or binding modes can be inferred from the structures, a strong binding mode mediated by stereospecific contacts between the MT and the whole KIF1A-MT interface, and a weak-binding mode mediated by flexible electrostatic interactions between the tubulin C-terminal tails, the KIF1A K-loop and possibly the neck-linker and the start of the coiled-coil domain. Regions of the structures with high mobility are illustrated with a blurred depiction. The PDB accession codes of the structure(s) representing each model step are displayed to the right, with different font colors indicating the KIF1A construct used, black for WT, red for the P305L mutant, and blue for the P364L mutant. The cycle starts with a KIF1A dimer in the ADP state interacting with the MT in a weakly bound mode (state 1). In this state, KIF1A could engage in biased one-dimensional diffusion (Supp. Fig 8) or in directed motion, where the two motor heads coordinate alternating conformations, nucleotide states, and MT binding modes to move in a hand-over-hand manner in the MT plus-end direction (steps 2 to 6). The P305L mutation alters the structure of the KIF1A MT interface, hindering the transition to the strongly bound configuration (from step 1 to step 2). Several points of the coordination between the two heads can be derived from the structures: In steps 2 and 3 one head is prevented from binding to the MT until the MT-bound head binds ATP (steps 4 and 5). This is because a two-heads-strongly-bound configuration cannot form unless one of the heads (the trailing head) has a docked neck linker, which occurs in the ATP-bound state. In step-4, the MT-bound head with ATP (head-1) has a reduced probability to reach the fully closed conformation until the partner motor domain binds strongly to the MT in the forward position (step-5). This is supported by the presence of distinct conformations of the trailing head in the ATP-bound state (see results). In the two-heads MT-bound configuration (step-5), inter-head tension and the differently oriented neck-linkers maintain the two heads in distinct conformations. In this configuration, nucleotide pocket closure and ATP hydrolysis in the leading head is paused until the trailing head detaches allowing the neck-linker of the leading head to dock. After ATP hydrolysis, product release and detachment of the rear head the neck-linker of the leading head docks moving the detached head forward (step-6). Steps 6 and 4 are equivalent but the two heads have interchanged leading and trailing positions and the KIF1A molecule is displaced 8 nm (the distance between tubulin heterodimers along a protofilament) in the MT + end direction.

The prevailing model attributes KIF1A’s superprocessivity primarily to the K-loop’s interaction with tubulin^82^, rather than tight coordination between the two motor domains. This perspective was partly informed by the assumption that KIF1A’s neck-linker is longer than that of kinesin-1^82,92^. Contrary to this, our structures reveal that the KIF1A neck-linker is actually shorter than that of kinesin-1 and other kinesins, suggesting a tighter connection between the motor domains when both are bound to the MT. Furthermore, our mutagenesis studies indicate that altering KIF1A’s neck-linker —by substituting the proline at the end of the neck-linker with the equivalent residue in kinesin-1— markedly reduces the motor’s run length. These results collectively favor a model where KIF1A’s superprocessivity is a synergistic outcome of enhanced MT binding via the K-loop and tight coordination between both motor domains.

Our model also revises traditional views of the mechanochemical cycle by introducing a state in which both motor domains are simultaneously ATP-bound while strongly bound to the MT (Fig. 7, step 5). This is contrary to common models suggesting that the leading head in the two-heads-bound configuration is nucleotide free, with ATP binding occurring only after the trailing head detaches from the MT^75,91,93^. However, our structural data lead us to dispute this concept. Firstly, we have observed a two-heads-bound configuration with nucleotides bound to both heads in KIF1A, a finding also corroborated by the high-resolution structures of KIF14^72^. Secondly, structural data do not support the restriction of ATP binding to the leading head, as the nucleotide-binding pocket exhibits an open conformation similar to that observed in the MT-bound Apo and ADP states (Fig. 2) ^72^. If ATP hydrolysis and product release occur rapidly in the trailing head^77^, the intermediate state where both heads are strongly bound to the MT (Fig. 7, step 5) would be short-lived. Consequently, the KIF1A dimer would spend a greater proportion of its translocation cycle in configurations where only one head is strongly bound to the MT.

Contrary to previous proposals suggesting that neck-linker docking in KIF1A and kinesin-1 occurs after ATP hydrolysis rather than upon ATP binding^77,94^, our data for KIF1A align with studies of other kinesins that associate neck-linker docking to ATP binding^72,95–97^. Intriguingly, in the ATP-bound state of KIF1A, we observed a significant proportion of molecules in a one-head-bound conformation with a docked neck-linker. This observation raises the possibility that ATP hydrolysis might facilitate the binding of the tethered head to the leading position, rather than being a prerequisite for neck-linker docking. However, our structural data also indicates that binding of the leading head is necessary for a more complete closure of the nucleotide-binding pocket of the trailing head (∼50% of the motors in the AMP-PNP one-head-bound configuration where in a semi-closed conformation, Fig. 2d, Supplementary Figs. 2f and 3). Assuming that the closed conformation is the catalytic one^97^, this would suggest that ATP hydrolysis is more favorable after the two-heads-bound configuration occurs.

Our data also elucidate the impact of the KAND mutation P305L on MT-binding (Fig. 7 step 1 to 2). Unexpectedly, the P305L mutation primarily influences the orientation of F303, rather than the adjacent highly conserved 3_10_-helix region of loop-12, which was previously shown to be critical for KIF1A-MT binding^84,98^. This altered orientation of F303 provides insights into the reduced MT-binding efficiency observed in the P305L mutant. Furthermore, our experiments reveal that the introduction of the F303V mutation alone reduced processivity, emphasizing the vital role of this residue in KIF1A’s motility. A similar modification of the corresponding residue in kinesin-1’s loop-12 also resulted in a reduced MT affinity^98^. Notably, combining the P305L and F303V mutations partially restored processivity, suggesting that this region of loop-12 may be a viable target for future structure-guided design strategies.

In summary, our findings support a revised model of the KIF1A mechanochemical cycle, where processivity is derived from enhanced K-loop-mediated interactions with MTs and from the coordination between the two motor domains. This revised model not only deepens our understanding of KIF1A’s superprocessivity but also sheds light on how mutations like P305L impact its function. Our structural insights into these mutations, especially their unexpected effects on MT binding, highlight potential areas for therapeutic interventions. These advancements in understanding the structural and functional aspects of KIF1A are pivotal for developing targeted treatments, particularly for KAND and related neurological disorders.

## Methods

### Generation of plasmids for KIF1A constructs

A plasmid for a previous published KIF1A construct^9^ (KIF1A(Homo sapiens, aa 1-393)-leucine zipper-SNAPf-EGFP-6His) was used as the template for all constructs in this study. For proteins used in the cryo-EM studies, the SNAPf-EGFP-6His tag was replaced with a strep-II tag (IBA Lifesciences GmbH) using Q5 mutagenesis (New England Biolabs Inc., #E0554S). Mutations within KIF1A were generated using Q5 mutagenesis. The K-loop swap mutant has EQANQRSV instead of the amino acid sequence of EMDSGPNKNKKKKKTD (KIF1A residues 287-302). All plasmids were confirmed by sequencing.

### Protein expression in *E. coli*

KF1A expression was performed as described in previous studies^9,52^. Briefly, each plasmid was transformed into BL21-CodonPlus(DE3)-RIPL competent cells (Agilent Technologies, #230280). A single colony was picked and inoculated in 1 mL of terrific broth (TB) (protocol adopted from Cold Spring Harbor protocol, DOI: 10.1101/pdb.rec8620) with 50 µg/mL carbenicillin and 50 µg/mL chloramphenicol. The 1-mL culture was shaken at 37 °C overnight, and then inoculated into 400 mL of TB (or 1–2 L for cryo-EM studies) with 2 µg/mL carbenicillin and 2 µg/mL chloramphenicol. The culture was shaken at 37 °C for 5 hours and then cooled on ice for 1 hour. IPTG was then added to the culture to a final concentration of 0.1 mM to induce expression. Afterwards, the culture was shaken at 16 °C overnight. The cells were harvested by centrifugation at 3,000 rcf for 10 minutes at 4°C. The supernatant was discarded, and 1.25 mL of B-PER™ Complete Bacterial Protein Extraction Reagent (ThermoFisher Scientific, #89821) per 100 mL culture with 2 mM MgCl_2_, 1 mM EGTA, 1 mM DTT, 0.1 mM ATP, and 2 mM PMSF was added to the cell pellet. The cells were fully resuspended, and flash frozen in liquid nitrogen. If the purification was not done on the same day, the frozen cells were stored at –80 °C.

### Protein purification

To purify the protein, the frozen cell pellet was thawed at 37 °C. The solution was nutated at room temperature for 20 minutes and then dounced for 10 strokes on ice to lyse the cells. Unless specified, the following procedures were done at 4 °C. The cell lysate was cleared by centrifugation at 80,000 rpm (260,000 rcf, *k*-factor=28) for 10 minutes in an TLA-110 rotor using a Beckman Tabletop Ultracentrifuge Unit. The supernatant was flown through 500 μL of Roche cOmplete™ His-Tag purification resin (Millipore Sigma, #5893682001) for His-tag tagged proteins, or 2 mL of Strep-Tactin® Sepharose® resin (IBA Lifesciences GmbH, #2-1201-002) for strep-II tagged proteins. The resin was washed with wash buffer (WB) (for His-tagged protein: 50 mM HEPES, 300 mM KCl, 2 mM MgCl_2_, 1 mM EGTA, 1 mM DTT, 1 mM PMSF, 0.1 mM ATP, 0.1% Pluronic F-127 (w/v), 10% glycerol, pH 7.2; for strep-II tagged protein, Pluronic F-127 and glycerol were omitted). For proteins with a SNAPf-tag, the resin was mixed with 10 μM SNAP-Cell® TMR-Star (New England Biolabs Inc., #S9105S) at room temperature for 10 minutes to label the SNAPf-tag. The resin was further washed with WB, and then eluted with elution buffer (EB) (for His-tagged protein: 50 mM HEPES, 150 mM KCl, 150 mM imidazole, 2 mM MgCl_2_, 1 mM EGTA, 1 mM DTT, 1 mM PMSF, 0.1 mM ATP, 0.1% Pluronic F-127 (w/v), 10% glycerol, pH 7.2; for strep-II tagged protein: 80 mM PIPES, 2 mM MgCl_2_, 1 mM EGTA, 1 mM DTT, 0.1 mM ATP, 5 mM desthiobiotin). The Ni-NTA elute was flash frozen and stored at –80 °C. The Strep-Tactin elute was concentrated using a Amicon Ultra-0.5 mL Centrifugal Filter Unit (30-kDa MWCO) (Millipore Sigma, #UFC503024). Storage buffer (SB) (80 mM PIPES, 2 mM MgCl_2_, 80% sucrose (w/v) was added to the protein solution to have a final 20% sucrose (w/v) concentration, and the protein solution was flash frozen and stored at −80 °C. The purity of the proteins was confirmed on polyacrylamide gels.

### Microtubule-binding and -release assay

An MT-binding and -release (MTBR) assay was performed to remove inactive motors for single-molecule TIRF assay. 50 μL of eluted protein was buffer-exchanged into a low salt buffer (30 mM HEPES, 50 mM KCl, 2 mM MgCl_2_, 1 mM EGTA, 1 mM DTT, 1 mM AMP-PNP, 10 µM taxol, 0.1% Pluronic F-127 (w/v), and 10% glycerol) using 0.5-mL Zeba™ spin desalting column (7-kDa MWCO) (ThermoFisher Scientific, #89882). The solution was warmed to room temperature and 5 μL of 5 mg/mL taxol-stabilized MTs was added. The solution was well mixed and incubated at room temperature for 2 minutes to allow motors to bind to the MTs and then spun through a 100 μL glycerol cushion (80 mM PIPES, 2 mM MgCl_2_, 1 mM EGTA, 1 mM DTT, 10 µM taxol, and 60% glycerol, pH 6.8) by centrifugation at 45,000 rpm (80,000 rcf, *k*-factor=33) for 10 minutes at room temperature in TLA-100 rotor using a Beckman Tabletop Ultracentrifuge Unit. Next, the supernatant was removed and the pellet was resuspended in 50 μL high salt release buffer (30 mM HEPES, 300 mM KCl, 2 mM MgCl_2_, 1 mM EGTA, 1 mM DTT, 10 μM taxol, 3 mM ATP, 0.1% Pluronic F-127 (w/v), and 10% glycerol). The MTs were then removed by centrifugation at 40,000 rpm (60,000 rcf, *k*-factor=41) for 5 minutes at room temperature. Finally, the supernatant containing the active motors was aliquoted, flash frozen in liquid nitrogen, and stored at –80 °C.

### Single-molecule TIRF motility assay

MTBR fractions were used for the single-molecule TIRF assay and the dilutions were adjusted to an appropriate density of motors on MTs. The assay was performed as described before^99^ except the motility buffer was BRB80 (80 mM PIPES, 2 mM MgCl_2_, 1 mM EGTA, 1 mM DTT, 10 µM taxol, 0.5% Pluronic F-127 (w/v), 2 mM ATP, 5 mg/mL BSA, 1 mg/mL α-casein, gloxy oxygen scavenging system, and 10% glycerol, pH 6.8). Images were acquired with 200 ms per frame (total 600 frames per movie), and then analyzed via a custom-written MATLAB software. Kymographs were generated using ImageJ2 (version 2.9.0). Statistical analysis was performed and graphs were generated using GraphPad Prism (version 9.5.0).

### Cryo-EM

#### Preparation of microtubules

Microtubules (MTs) were prepared from porcine brain tubulin (Cytoskeleton, Inc. CO). Tubulin lyophilized pellets were resuspended in BRB80 (80 mM K-PIPES, 1 mM MgCl_2_, 1 mM EGTA, pH 6.8) to 5 mg/mL and spun at 313,000 × *g* before polymerization to eliminate aggregates. MT polymerization was done in conditions to enrich the number of MTs with 15 protofilaments^100^ as follows. The clarified resuspended tubulin solution was supplemented with 2 mM GTP, 4 mM MgCl_2_, 12%(*v*/*v*) DMSO and incubated 40 minutes at 37 °C. An aliquot of stock Paclitaxel (Taxol^®^) solution (2 mM in DMSO) was added for a final paclitaxel concentration of 250 µM and incubated for another 40 minutes at 37 °C. The MTs were then spun at 15,500 × *g*, 25 °C and the pellet resuspended in BRB80 with 20 μM paclitaxel.

#### Cryo-EM of KIF1A-MT complexes

Four µL of 6 μM MT solution in BRB80 plus 20 μM paclitaxel were layered onto (UltrAuFoil R1.2/1.3 300 mesh) plasma cleaned just before use (Gatan Solarus plasma cleaner, at 15 W for 6 seconds in a 75% argon/25% oxygen atmosphere), the MTs were incubated 1 minute at room temperature and then the excess liquid removed from the grid using a Whatman #1 paper. Four µL of a solution of KIF1A WT in BRB80 supplemented with 20 μM paclitaxel (kinesin concentrations given in Supplementary Table 1) and either (1) 4 mM AMP-PNP, (2) 4 mM ADP, or (3) 5 × 10^−3^ units per µL apyrase (APO condition) were then applied onto the EM grid and incubated for 1 minute at room temperature. The grid was mounted into a Vitrobot apparatus (FEI-ThermoFisher MA), incubated 1 minute at room temperature and plunge-frozen into liquid ethane (Vitrobot settings: 100% humidity, 3 seconds blotting with Whatman #1 paper and 0 mm offset). Grids were stored in liquid nitrogen. For the P305L mutant, the kinesin solution was prepared as the solution for WT KIF1A with the difference that instead of BRB80 we used BRB36 (36 mM K-PIPES, 1 mM MgCl_2_, 1 mM EGTA, pH 6.8) (Supplementary Table 1).

#### Cryo-EM data collection

Data were collected at 300 kV on Titan Krios microscopes equipped with a K3 summit detector. Acquisition was controlled using Leginon^101^ with the image-shift protocol and partial correction for coma induced by beam tilt^102^. The pixel sizes, defocus ranges and cumulated dose are given in Table 1.

#### Processing of the cryo-EM datasets of MT-kinesin complexes

The processing was done similar as previously described^72^. Movie frames were aligned with Relion generating both dose-weighted and non-dose-weighted sums. Contrast transfer function (CTF) parameters per micrographs were estimated with Gctf^103^.

Helical reconstruction on 15R MTs was performed using a helical-single-particle 3D analysis workflow in Frealign^104^, as described previously^72,88^ with 664 pixels box size, with each filament contributing only to one of the two half dataset. Per-particle CTF refinement was performed with FrealignX^105^.

To select for tubulins bound to kinesin motors and to improve the resolution of the kinesin-tubulin complexes, the procedure HSARC^72^ was used for these one-head-bound configurations. The procedure follows these steps:

1. Relion helical refinement. The two independent Frealign helical refined half datasets were subjected to a single helical refinement in Relion 3.1^106^ where each microtubule was assigned to a distinct half-set and using as priors the Euler angle values determined in the helical-single-particle 3D reconstruction (initial resolution: 8 Å, sigma on Euler angles sigma_ang: 1.5, no helical parameter search).
2. Asymmetric refinement with partial signal subtraction. Atomic models of kinesin-tubulin complexes derived from our recent work on KIF14^72^ were used to generate a soft mask mask_full_ using EMAN pdb2mrc and relion_mask_create (low-pass filtration: 30 Å, initial threshold: 0.05, extension: 14 pixels, soft edge: 8 pixels). For the one-head-bound only datasets (MT-KIF1A-ADP, MT-KIF1A-APO, MT-KIF1A^P305L^-ADP and MT-KIF1A^P305L^-APO), mask_full_ was generated with a model containing 1 kinesin motor bound to 1 tubulin dimer and two longitudinally flanking tubulin subunits. For the datasets containing two-heads-bound configurations (MT-KIF1A-ANP, MT-KIF1A^P364L^-ANP and MT-KIF1A^P305L^-ANP), mask_full_ was generated from a kinesin dimer model bound to two tubulin dimers. The helical dataset alignment file was symmetry expanded using the 15R MT symmetry of the dataset. Partial signal subtraction was then performed using mask_full_ to retain the signal within that mask. During this procedure, images were re-centered on the projections of 3D coordinates of the center of mass of mask_full_ (C_M_) using a 416 pixels box size. The partially signal subtracted dataset was then used in a Relion 3D refinement procedure using as priors the Euler angle values determined form the Relion helical refinement and the symmetry expansion procedure (initial resolution: 8 Å, sigma_ang: 5, offset range corresponding to 5 Å, healpix_order and auto_local_healpix_order set to 5). The CTF of each particle was corrected to account for their different position along the optical axis.
3. 3D classification of the kinesin signal. A mask mask_kinesin_ was generated like in step (2) but using only the kinesin coordinates (a single kinesin head for the datasets MT-KIF1A-ADP, MT-KIF1A-APO, MT-KIF1A^P305L^-ADP and MT-KIF1A^P305L^-APO; a two-heads-bound kinesin dimer for the dataset MT-KIF1A-ANP, MT-KIF1A^P364L^-ANP and MT-KIF1A^P305L^-ANP). A second partial signal subtraction procedure identical to first one in step (2) but using mask_kinesin_ instead of mask_full,_ with particles re-centered on the projections of C_M_ was performed to subtract all but the kinesin signal. The images obtained were resampled to 3.5 Å/pixel in 100-pixel boxes and the 3D refinement from step 2 was used to update the Euler angles and shifts of all particles. For the datasets MT-KIF1A-ADP, MT-KIF1A-APO, MT-KIF1A^P305L^-ADP and MT-KIF1A^P305L^-APO with a single-head-bound configuration, a 3D focused classification without images alignment and using a mask for the kinesin generated like mask_kinesin_ (low-pass filtration: 30 Å, initial threshold: 0.9, extension: 1 pixel, soft edge: 3 pixels) was then performed on the resampled dataset (8 classes, tau2_fudge: 4, padding: 2, iterations: 175). The class(es) showing a kinesin were selected and merged for the next step. For the MT-KIF1A^P305L^-ADP which had very low decoration (∼9 %, Supplementary Table 1), the previous classification led to a single decorated class (14% of the dataset) with a weak kinesin signal. Particles from this class were further classified in a second 3D classification (4 classes, tau2_fudge: 4, padding: 2, iterations: 175) and the main decorated class (68%) with a recognizable kinesin was selected while the others (with resolution lower than 15 Å) were not. For the MT-KIF1A-ANP, MT-KIF1A^P364L^-ANP and MT-KIF1A^P305L^-ANP datasets, 3D classifications in 8 classes performed as described above on the partially subtracted data with 2 kinesin motor domains left revealed that both one-head- and two-heads-bound configurations were present but not fully separated. The following hierarchical 3D-classification was used instead (Supplementary Figs. 2, 3 and 9). Two different masks were generated (Supplementary Figure 2a-c): one covering the kinesin site closest to the (-) end of the MT called mask_T_, and the other covering the kinesin site closest to the (+) end of the MT, called mask_L_. These two masks cover respectively the trailing head and leading head of a two-heads-bound kinesin dimer bound to these two sites (sites T and L respectively, Supplementary Figure 2a-c). Each of these two masks was generated from an atomic model corresponding to the overlay of a trailing head and a leading head of a two-heads-bound kinesin dimer with its associated coiled-coil, so that the focused classification includes the signal of the coiled-coil independent of the kinesin registration (Supplementary Figure 2d-e). These models were converted to maps with EMAN pdb2mrc and the masks were generated with relion_mask_create (low-pass filtration: 30 Å, initial threshold: 1.3, extension: 1 pixel, soft edge: 3 pixels). First a 3D classification was performed focusing the classification on mask_T_ (8 classes, tau2_fudge: 6, padding: 2, iterations: 175, Supplementary Figure 2f). Then a second focused classification using mask_L_ (4 classes, padding: 2, iterations: 175) was performed on the classes obtained in the previous classification step and selecting only the classes that were potentially trailing heads of a two-heads-bound configuration (Supplementary Figure 2f). This excluded classes that contained clear neck-linker and coiled-coil cryo-EM densities toward the (-) end of the motor-domain density (i.e. a leading head), the classes that show no kinesin density and the classes having an unrecognizable signal. Each of the main classes used was named as indicated in Supplementary Figure 2f. The decoration and propensity of the states identified resulting from these classifications are given in Supplementary Table 1. All datasets produced at least one class where the kinesin motor densities were well-resolved.
4. 3D reconstructions with original images (not signal subtracted). To avoid potential artifacts introduced by the signal subtraction procedure, final 3D reconstructions of each half dataset were obtained using relion_reconstruct on the original image-particles extracted from the micrographs without signal subtraction. To increase the resolution on the site T for the datasets MT-KIF1A-ANP and MT-KIF1A^P305L^-ANP, some reconstructions were generated by merging the data from different classes obtained in the classification on the site T or L. In this case, the names of the classes are concatenated, for example, MT-KIF1A-ANP-T_23_L_1_ was generated by merging the data from MT-KIF1A-ANP-T_2_L_1_ and MT-KIF1A-ANP-T_3_L_1_.
5. To obtain a final locally filtered and locally sharpened map for the states listed in table 1 (MT-KIF1A-ANP-T_23_L_1_, MT-KIF1A-ANP-T_2_L_1_, MT-KIF1A-ANP-T_3_L_1_, MT-KIF1A-ANP-T_1_L_02*_, MT-KIF1A-ADP, MT-KIF1A-APO, MT-KIF1A^P364L^-ANP, MT-KIF1A^P305L^-ANP-TL_1_, MT-KIF1A^P305L^-ANP-TL_012*_, MT-KIF1A^P305L^-ADP and MT-KIF1A^P305L^-APO), post-processing of the pair of unfiltered and unsharpened half maps was performed as follows. One of the two unfiltered half-map was low-pass-filtered to 15 Å and the minimal threshold value that does not show noise around the MT fragment was used to generate a mask with relion_mask_create (low-pass filtration: 15 Å, extension: 10 pixels, soft edge: 10 pixels). This soft mask was used in blocres^107^ on 12-pixel size boxes to obtain initial local resolution estimates. The merged map was locally filtered by blocfilt^107^ using blocres local resolution estimates and then used for local low-pass filtration and local sharpening in localdeblur^108^ with resolution search up to 25 Å. The localdeblur program converged to a filtration appropriate for the tubulin part of the map but over-sharpened for the kinesin part. The maps at every localdeblur cycle were saved and the map with better filtration for the kinesin part area was selected with the aim to improve the resolution of the kinesin loops.

The final reconstructions of MT-KIF1A-ANP-T_1_L_1_, MT-KIF1A-ANP-T_1_L_0_, MT-KIF1A-ANP-T_2_L_0_, MT-KIF1A-ANP-T_3_L_0_, MT-KIF1A-ANP-T_1_L_*_, MT-KIF1A-ANP-T_2_L_*_, MT-KIF1A-ANP-T_3_L_*_ (Supplementary Fig.3) were low-passed filtered at 4.0 Å and sharpened with a b-factor of −40 A^2^.

#### Cryo-EM resolution estimation

The final resolutions for each cryo-EM class average reconstruction listed in table 1 (MT-KIF1A-ANP-T_23_L_1_, MT-KIF1A-ANP-T_2_L_1_, MT-KIF1A-ANP-T_3_L_1_, MT-KIF1A-ANP-T_1_L_02*_, MT-KIF1A-ADP, MT-KIF1A-APO, MT-KIF1A^P364L^-ANP MT-KIF1A^P305L^-ANP-TL_1_, MT-KIF1A^P305L^-ANP-TL_012*_, MT-KIF1A^P305L^-ADP and MT-KIF1A^P305L^-APO) were estimated from FSC curves generated with Relion 3.1 postprocess (FSC_0.143_ criteria, Supplementary Figure 4). To estimate the overall resolution, these curves were computed from the two independently refined half maps (gold standard) using soft masks that isolate a single asymmetric unit containing a kinesin and a tubulin dimer. The soft masks were created with Relion 3.1 relion_mask_create (for MT datasets: low pass filtration: 15 Å, threshold: 0.1, extension: 2 pixels, soft edge: 5 pixels) applied on the correctly positioned EMAN pdb2mrc density map generated with the coordinates of the respective refined atomic models. FSC curves for the tubulin or kinesin parts of the maps were generated similarly using the corresponding subset of the PDB model to mask only a kinesin or a tubulin dimer (Supplementary Fig. 4, Table 1).

The final cryo-EM maps together with the corresponding half maps, the masks used for resolution estimation, the masks used in the partial signal subtraction for the MT datasets, the low-passed filter maps used in Fig. 5 and Supplementary Figs. 6 and 7, and the FSC curves (Supplementary Fig.4) are deposited in the Electron Microscopy Data Bank (Table 1).

### Model building

Atomic models of the cryo-EM density maps were built as follows. First, atomic models for each protein chains were generated from their amino-acid sequence by homology modeling using Modeller^109^. Second, the protein chains were manually placed into the cryo-EM densities and fitted as rigid bodies using UCSF-Chimera^110^. Third, the models were flexibly fitted into the density maps using Rosetta for cryo-EM relax protocols^111,112^ and the models with the best scores (best match to the cryo-EM density and best molprobity scores) were selected. Fourth, the Rosetta-refined models were further refined against the cryo-EM density maps using Phenix real space refinement tools^113^. Fifth, the Phenix-refined models were edited using Coot^114^. Several iterations of Phenix real space refinement and Coot editing were performed to reach the final atomic models. Atomic models and cryo-EM map figures were prepared with UCSF-Chimera^110^ or USCF ChimeraX^115^ and R^116^.

### Nucleotide binding pocket openness

Nucleotide binding pocket (NP) openness values for each atomic model were calculated as the average of six distances between KIF1A Cα carbons of highly conserved residues (R216 and A250 to P14, to S104 and to Y105) across the kinesin nucleotide-binding pocket^74^. Atom distances were calculated from the atomic models using UCSF-Chimera^110^. Precision of the NP openness values (expressed as mean ± standard deviation (SD)) was estimated by regenerating multiple atomic models^117^. For each atomic model eighty models were regenerated by flexibly fitting the model into the corresponding cryo-EM map using Rosetta for cryo-EM relax protocols^111,112,117^. Mean and SD were calculated for each model from six NP openness values: one from the model itself and the others from the five best-scored Rosetta-regenerated models.

## Acknowledgements

This work was supported by National Institutes of Health Grant R01GM113164 (H.S.), R01GM147332 (A.G. and H.S.), R01GM098469 (A.G.), and R01NS114636 (A.G.). Cryo-EM data collection was performed at the Simons Electron Microscopy Center and National Resource for Automated Molecular Microscopy located at the New York Structural Biology Center, supported by grants from the Simons Foundation (SF349247), NYSTAR, and the NIH National Institute of General Medical Sciences (GM103310) with additional support from Agouron Institute (F00316) and NIH (OD019994).

## Author contributions

L.R. generated, expressed, and purified the constructs for the cryo-EM and single-molecule studies. L.R. performed single-molecule experiments, collected, and analyzed the data, and interpreted the results. M.P.M.H.B. and A.B.A. assembled kinesin-MT complexes and made cryo-EM grid samples; A.B.A. performed sample screening and optimization for cryo-EM imaging and performed MT selection; M.P.M.H.B., A.B.A. and H.S. designed the cryo-EM experiments; M.P.M.H.B. and A.B.A. performed cryo-EM data collection; M.P.M.H.B. designed and performed the cryo-EM data processing of kinesin-MT complexes and interpreted the results; M.P.M.H.B. and H.S. built and refined the atomic models, and interpreted the structures; A.G. and H.S. conceived and coordinated the project and interpreted the results; M.P.M.H.B., L.R, A.B.A., A.G., and H.S. wrote the manuscript.

## Supplementary Figures and Tables

**Supplementary Fig 1.**
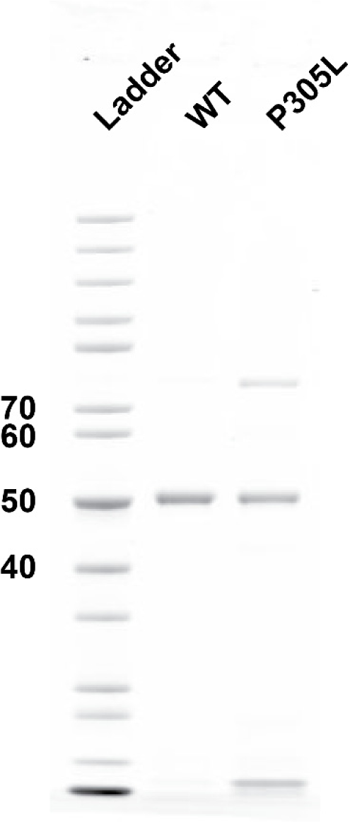
Polyacrylamide gel of KIF1A constructs. Coomassie blue-stained gel of KIF1A (KIF1A(aa1-393)-LZ-strepII) (WT) and KIF1A^P305L^(KIF1A(aa1-393, P305L)-LZ-strepII) (P305L); Ladder: molecular weight standards. The molecular weight of the ladder is indicated on the left in kDa.

**Supplementary Fig 2.**
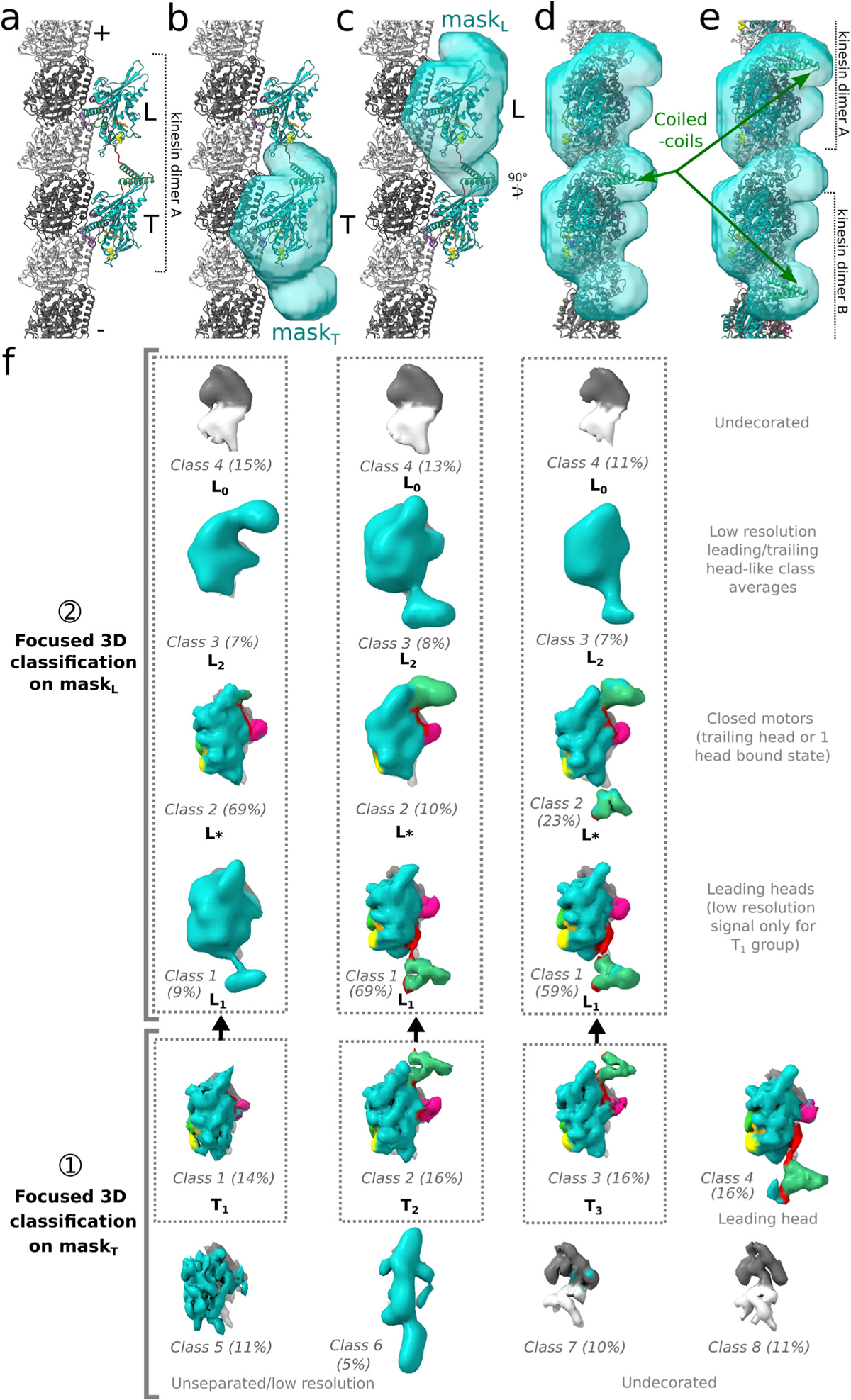
3D-classification strategy used for the MT-KIF1A-ANP dataset. **a:** Model of a KIF1A kinesin dimer (“kinesin dimer A”) on a microtubule protofilament. The polarity of the microtubule is indicated. The leading head of the kinesin dimer is indicated by the letter L while the trailing head is indicated by the letter T. Coiled-coil is colored green and other colors are as in Fig.1. **b-c:** Semi-transparent surface view of the two masks mask_T_ (**b**) and mask_L_ (**c**) used successively during the 3D classification. These two 30 Å-resolution masks differ only by their position: mask_T_ covers the trailing position of the kinesin dimer A (site T, **b**), while mask_L_ covers the leading position (site L, **c**) at the next kinesin-binding site toward the (+) end of the microtubule protofilament. **d:** 90 degree rotated view of the kinesin dimer model of panel (**a**) with both mask_T_ and mask_L_ displayed. **e:** Same view as (**d**) with 2 consecutive two-heads-bound kinesin dimers A and B attached to the microtubule protofilament but with a registration off by 82 Å (length of tubulin dimer) compared to the kinesin dimer A shown in (**a-e**). Note that both masks cover the area occupied by the coiled-coil densities for the two types of registrations displayed in (**e**) and (**d**). **f:** Scheme representing the classification strategy used for the MT-KIF1A-ANP dataset using the mask_T_ and mask_L_ illustrated in (**a-e**). All class averages shown in the panel correspond to the 3.5Å/pixel class averages produced by the 3D classifications, maintaining the same viewing orientation as panels (**d-e**). In the initial step (step➀), a 3D classification in 8 classes and focusing on mask_T_ was performed. This led to the 8 class averages displayed on the lower part of the figure. The classes 1-3 correspond to kinesin motors in closed conformationss with the neck-linker docked, obtained in approximately equal proportions (refer to Supplementary Fig. 3 for a full-resolution side view of these three classes). These classes haven been designated as T_1_, T_2_ and T_3_. Class 4 corresponds to a leading head configuration, featuring an undocked neck-linker pointing backwards and connected to a coiled-coil structure. Class 5 contains a kinesin-like density but its specific state could not be confidentially assigned. It likely represents particles for which the state has not been effectively separated, unlike those classes 1 to 4. Classes 6 corresponds to a low-resolution class which an unassignable state. In contrast, classes 7 and 8 are devoid of any distinct decoration. Due to the symmetry expansion employed, the leading heads observed in the two-heads-bound configuration, as exemplified by class 4, are also expected to be present on site L. Given this, and considering that only the classes T_1_, T_2_ and T_3_ could correspond to two-heads-bound configuration kinesin dimers with their leading head on site L, these three classes were further classified on site L. In the second classification step (step ➁), with a focus on mask_L_, the particles from each of the T_1_, T_2_ and T_3_ classes were further divided into four subclasses. The corresponding class averages are presented in the figure. These class averages can be grouped into four distinct categories, as indicated in the figure, with some differences in relative frequencies. Each of T_1_, T_2_ and T_3_ has a major class at position L, containing respectively 69%, 69% and 59% of their particles. This major class, denoted as ‘L_*_‘, represents a motor in a closed conformation (i.e., belonging to another dimer) for T_1_ (specifically, class-2 in T_1_ classification at site L with mask_L_). It is worth noting that unless the kinesin coiled-coil were to unfold, the motor observed on the site L must correspond to another kinesin molecule. This indicates that the kinesin motor on site T, as seen in both class T_1_ and sub-class L_*_ (abbreviated T_1_L_*_), exists in a one-head-bound configuration. In contrast to T_1_, for T_2_ and T_3_, the major class observed on site L represents a connected leading head that is part of a two-heads-bound kinesin configuration. These classes, characterized by connected leading heads, were designated as ‘L_1_’. In the T_1_ classification on site L, only a weak and low-resolution class exhibited similarity to a leading head (Class 1, named L_1_; see also Supplementary Fig. 3 for a full-size reconstruction of this class), so T_1_L_1_ does not appear to be a pure two-heads-bound configuration class, unlike T_2_L_1_ and T_3_L_1_. Each of the classifications on site L for T_2_ and T_3_ also includes a class (class 2) corresponding to motors in a closed conformation with a coiled-coil density towards the (+) end of the motor (see Supplementary Fig. 3). Similar to the T_1_ classification, these classes were named L_*_ and they indicate the presence of one-head-bound configurations on the T site. In addition, each classifications on site L from T_1_, T_2_ and T_3_, yielded other low-resolution classes resembling either leading or trailing-head-like classes (classes 3, named L_2_), as well as empty sites (classes 4, named L_0_) without kinesin densities. The presence of these empty sites on site L indicates that the corresponding motors on site T (classes T_1_L_0_, T_2_L_0_ and T_3_L_0_) are in a one-head-bound configuration.

**Supplementary Fig 3.**
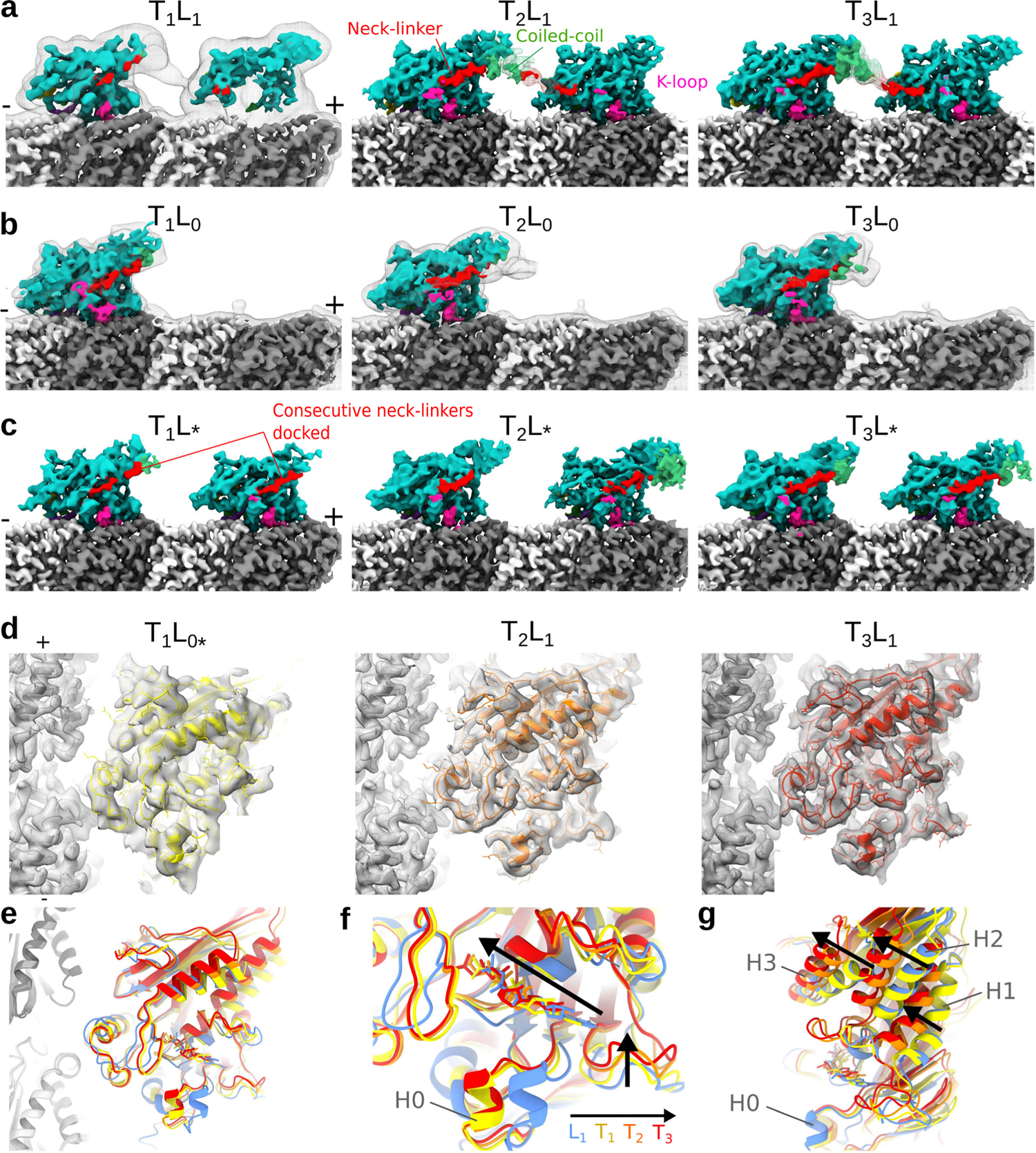
Multiple binding modes and conformational states of KIF1A-ANP. **a-c:** Isosurface representations of the cryo-EM maps from the KIF1A-ANP dataset for the classes T_1_L_1_, T_2_L_1_ and T_3_L_1_ (**a**), T_1_L_0_, T_2_L_0_ and T_3_L_0_ (**b**), T_1_L_*_, T_2_L_*_ and T_3_L_*._ **c**: The color scheme is the same as the one used in Fig. 1. The polarities of the microtubule protofilaments are indicated. To visualize the mobile K-loop region better, the signal around it was low-passed filtered and displayed at a lower contour level compared to the rest of the map. It’s important to highlight that in (**a**), the motor domain at site L exhibits a weaker signal compared to the one at site T. Only through low-pass filtering of this class average, as illustrated by the gray mesh, was it possible to detect a weak signal of connectivity between the two heads. Therefore, the small T_1_L_1_ class, accounting for 9% of the site L classification within the T_1_ class, appears to be a heterogeneous class that cannot be confidently assigned as being a two-heads-bound configuration. Since the other T_1_ classes are in a one-head-bound configuration, the vast majority of the T1 motors are therefore in a one-head-bound configuration, unlike T_2_ and T_3_. In T_2_L_1_ and T_3_L_1_, the coiled-coil density and part of the neck-linker from the leading head were low-pass filtered and displayed as a mesh at a lower contour level than the main map. Both T_2_L_1_ and T_3_L_1_ represent two-heads-bound configurations, as evidenced by the densities of the coiled-coil and the neck-linkers positioned in opposite orientations. Notably, there is an increasing density of the inter-head connection (neck-linker and coiled-coil) from T_1_L_1_ to T_2_L_1_ to T_3_L_1_, in these classes, accounting for 1.3%, 11% and 10% of the particles in the MT-KIF1A-ANP dataset, respectively. The T_3_L_1_ class exhibits the strongest inter-head connection density (neck-linker and coiled-coil) in the entire MT-KIF1A-ANP dataset, despite having fewer particles than T_2_L_1_, suggesting a more rigid conformation than T_2_L_1_. In (**b**), the densities of T_1_L_0_, T_2_L_0_ and T_3_L_0_ are displayed, overlaid with low-pass filtered versions of these maps. These maps show a lack of kinesin density at site L, indicating that the motors detected on the site T are in a one-head-bound configuration. In (**c**), each of the T_1_L_*_, T_2_L_*_ and T_3_L_*_ classes exhibits a docked neck-linker in both sites T and L, indicating that the motors detected on the site T are in a one-head-bound configuration, and that the motor seen on the site L corresponds to another kinesin dimer. **d-g:** Comparison of the conformations of the three detected KIF1A trailing heads, T_1_, T_2_ and T_3_. For this comparison, maps of the one-head-bound-configuration T_1_L_0*_, and of the two-heads-bound configurations T_2_L_1_ and T_3_L_1_ were used. In (**d**), semi-transparent isosurfaces of the nucleotide-binding area of the kinesin are shown, with underlying models displayed with a cartoon representation. T_1_, T_2_ and T_3_ models are overlaid with the same viewing angle in (**e**) and colored yellow, orange and red, respectively, as indicated. Additionally, the model of the leading head L_1_ (from T_23_L_1_) is displayed in blue for comparison. T_1_, T_2_ and T_3_ share the same H0 position, which is distinct from the open conformation found in L_1_. Notably, the positions of H2 close to the nucleotide and loop-5 (L5) follow the order L_1_, T_1_, T_2_, T_3_, as indicated by the black arrow. This trend corresponds to a progressive closing of the nucleotide-binding pocket, with the nucleotide inserted further in the pocket. This gradient of conformation is particularly visible in the helices H1 and H2. Panels (**f-g**) provide a similar comparison as in (**d-e**), focusing on these two helices. Arrows on the model overlay in (**g**) emphasize the gradation of motor closing from L_1_ to T_3_, with T_2_ and T_3_ being the most similar among these four conformations. T_1_ represents a semi-closed conformation (Fig. 2d, Supplementary Fig. 5) with both a docked neck-linker (**b-c**) and areas where it appears more similar to an open conformation (L_1_) than T_2_ and T_3_, such as the location of helices H1, H2 and H3. Importantly, T_1_ is a major conformation of the MT-bound head of KIF1A in the one-head-bound configuration:12% of the dataset in T_1_ (T_1_L_0*_) vs. 4% for T_2_ (T_2_L_0*_) and 6% for T_3_ (T_2_L_0*_) are in one-head-bound configurations, as shown in Supplementary Fig. 2. However, it has only a very low representation in the two-heads-bound configuration (< 1.3% (T_1_L_1_) vs. 11% for T_2_ (T_2_L_1_) and 10% for T_3_ (T_3_L_1_). Therefore, these structural observations indicate that the binding of the leading head restricts the conformation of the trailing head to a more closed conformation (T_2_ and T_3_), which is presumably more favorable for ATP hydrolysis. This property of KIF1A, to have the dynamic nucleotide-binding pocket of the MT-bound head partially closed in the one-head-bound configuration and further closed once the leading head binds, is possibly related to its high processivity.

**Supplementary Fig 4.**
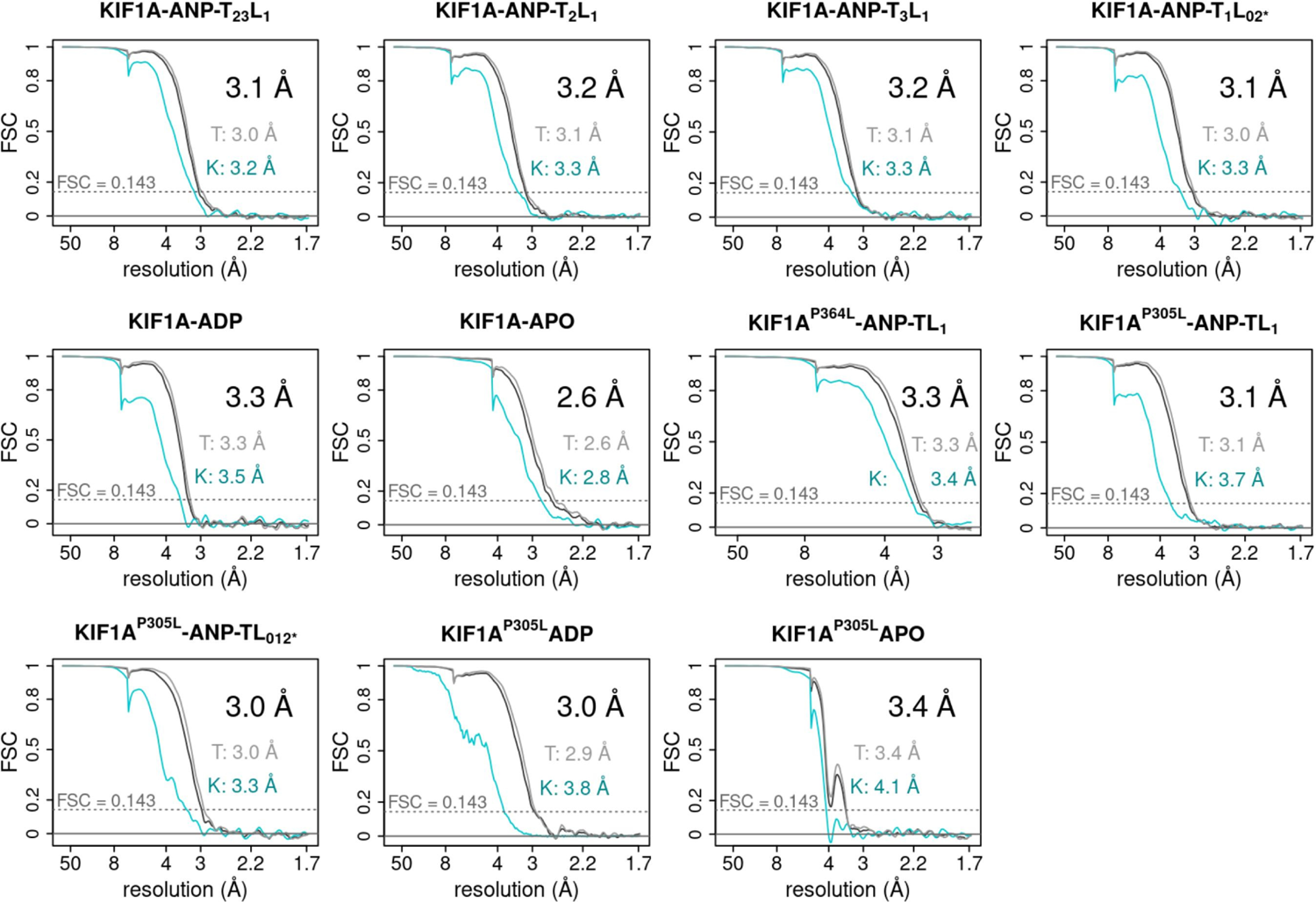
Resolution estimation of the cryo-EM maps. The figure displays FSC curves for KIF1A structures determined by cryo-EM. The overall resolution is represented by the black curve, the resolution for the tubulin (T) component is shown in grey, and the resolution for the kinesin (K) component is depicted in turquoise. The resolution values (FSC_0.143_) for the overall structure, the tubulin part, and the kinesin part are indicated. The half maps and masks used to generate the FSC curves have been deposited in the EMDB, with accession numbers provided in Table 1.

**Supplementary Fig 5.**
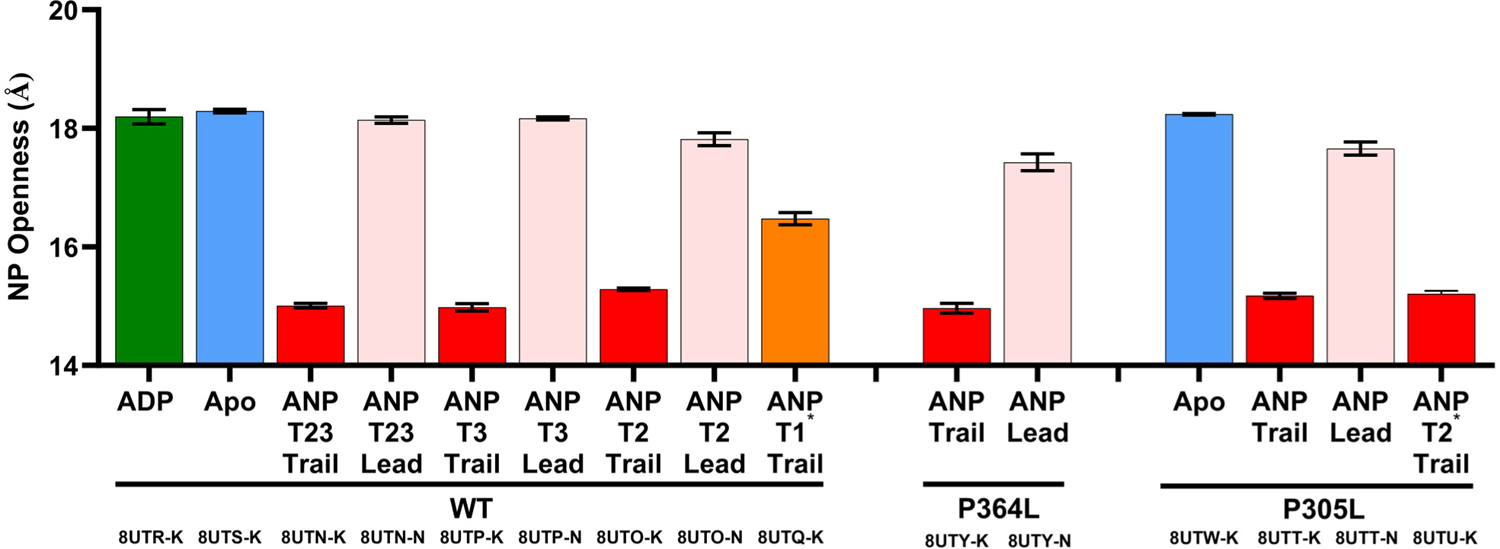
Nucleotide Pocket (NP) Openness. NP openness was estimated as the average of six distances between Cα carbons of selected residues (R216 and A250 to P14, to S104, and to Y105) across the KIF1A nucleotide-binding pocket. Each column bar corresponds to an atomic model of the KIF1A motor domain. The PDB accession code and the chain of the KIF1A motor domain are indicated at the bottom of the figure (PDB accession code-chain). The NP openness was not calculated for the P305L-ADP model (8UTV-K) due to the lower resolution of the kinesin motor domain in the corresponding cryo-EM map (Table 1), rendering this calculation unreliable. Column height and error bars correspond respectively to the mean and standard deviation of six openness values. These six openness values for each model correspond to the one calculated from the given model and the ones calculated from the best five of eighty regenerated Rosetta models (see methods).

**Supplementary Fig 6.**
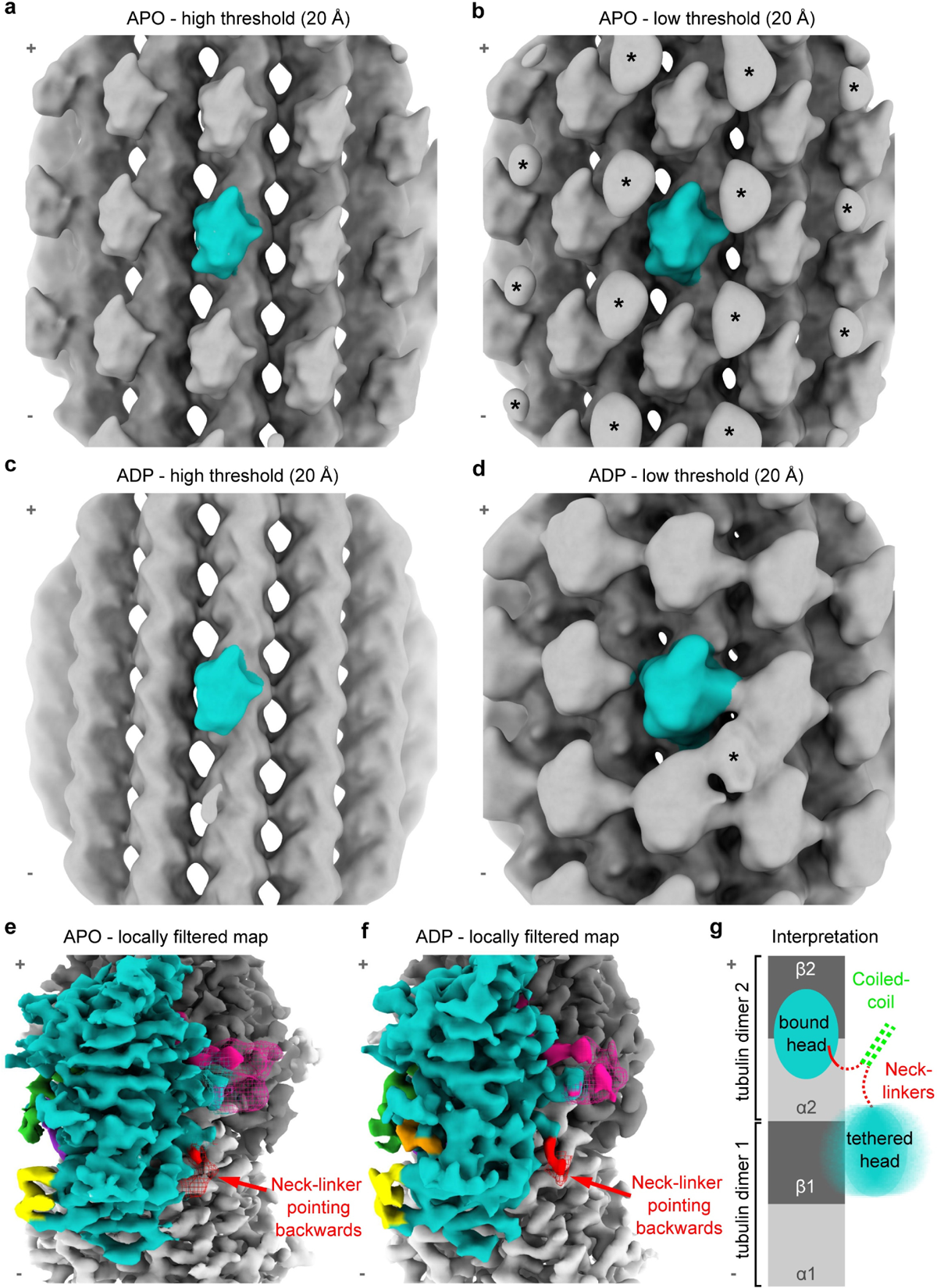
Tethered Head Location. **a-b:** Shown are 20 Å low-passed filtered cryo-EM maps of WT KIF1A in the Apo state (MT-KIF1A-APO) at high and low threshold, respectively. The kinesin belonging to the central asymmetric unit, on which the refinement was focused, is colored blue. Notably, in (b), extra densities indicated with an asterisk (*) are visible between the MT-bound kinesin motors and away from the MT surface. Such densities were not observed in the ANP dataset. All the 20 Å low-passed filtered cryo-EM maps displayed in this figure have been deposited as additional maps in the corresponding EMDB entries. **c-d:** Similar to (**a-b**), but for WT KIF1A in the ADP state (MT-KIF1A-ADP). Due to the lower decoration (approximately 32%, as shown in Supplementary Table 1) of this dataset (**c**), only one extra density (*) is detected near the central kinesin motor (**d**). This suggests that this density is associated with the central motor and that the similar extra densities seen in (**b**) are also linked to the nearest motor on the (+) end of the same protofilament. These extra densities could correspond to the position of the mobile unbound head or possibly to the coiled-coil region. In either case, this indicates that the associated tethered head is positioned backwards. **e-f:** Isosurfaces of the maps of WT KIF1A in the Apo state (MT-KIF1A-APO) (**e**) and WT KIF1A in the ADP state (MT-KIF1A-ADP) (**f**). Densities for the weaker K-loop and neck-linker were low-passed filtered and are represented as a mesh. Note that in both cases, the first residues of the neck-linker are pointing backward, towards the (-) end of the MT. **g:** Proposed interpretation of the combined observations in (**a-f)** for KIF1A in ADP and APO states. The resolved initial segment of the neck-linker and the extra low-resolution density present toward the (-) end of the motor suggests that the tethered head samples at least a position where it is oriented backward from the MT-bound motor.

**Supplementary Fig 7.**
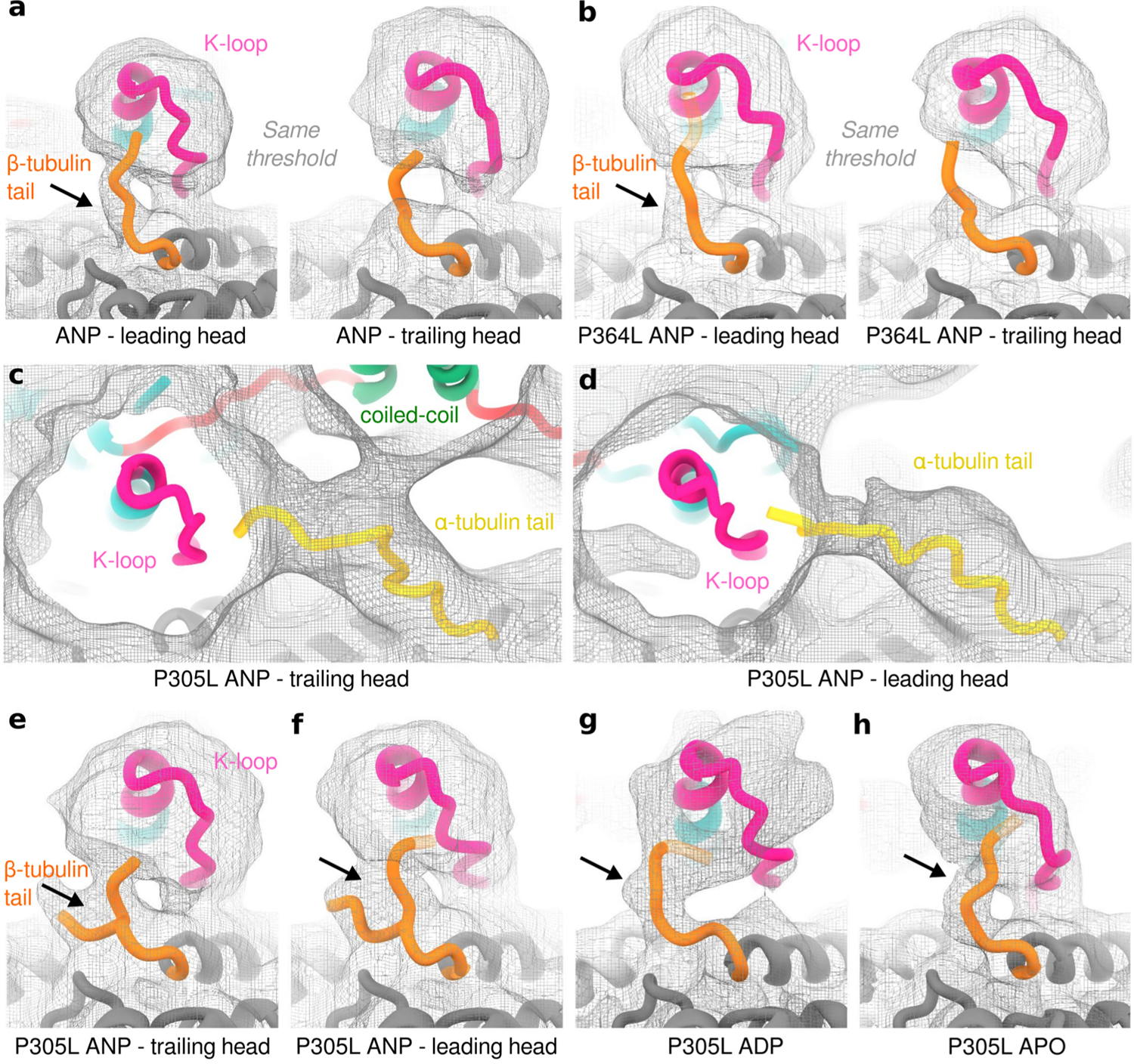
Interaction of the K-loop with the C-terminal tubulin tails. **a:** Shown is a 6 Å low-passed filtered cryo-EM map of the WT KIF1A two-heads-bound configuration in the ANP state (MT-KIF1A-ANP-T_23_L_1_), highlighting the areas around the β-tubulin tails near the leading and trailing heads of KIF1A, displayed at the same threshold. Notably, the density assigned to the β-tubulin tail is more resolved for the tail near the leading head than for the one near the trailing head of KIF1A. **b:** Similar to (**a**) but for the KIF1A P364L mutant two-heads-bound configuration in the ANP state (MT-KIF1A^P364L^-ANP-TL_1_). **c-d:** Displayed are 8 Å low-passed filtered cryo-EM map of the two-heads-bound configuration of the KIF1A P305L mutant (KIF1A^P305L^-ANP-TL_1_), showing densities of α-tubulin C-terminal tails reaching the K-loops of the trailing (**c**) or leading (**d**) head. Similar to WT KIF1A (Fig. 5b, c), the α-tubulin tail interacting with the trailing head is located within a pocket of positively-charged residues due to the charges of K-loop, the nearby coiled-coil, and the docked neck-linker. Given the shape of the map in this area, it is likely that additional interactions between the α-tubulin tail and the KIF1A coiled-coil occur. **e-h:** Presented are 6 Å low-passed filtered maps of KIF1A near the K-loop in distinct nucleotide states and motor-domain conformations. The density associated with the β-tubulin C-terminal tail is indicated with an arrow, and the threshold was adjusted for each map. It is worth noting that in the ANP state, the β-tubulin tail density is better resolved in the leading head than in the trailing head. The 6 Å and 8Å low-passed filtered maps used in this figure have been deposited as additional maps in the corresponding EMDB entries.

**Supplementary Fig. 8.**
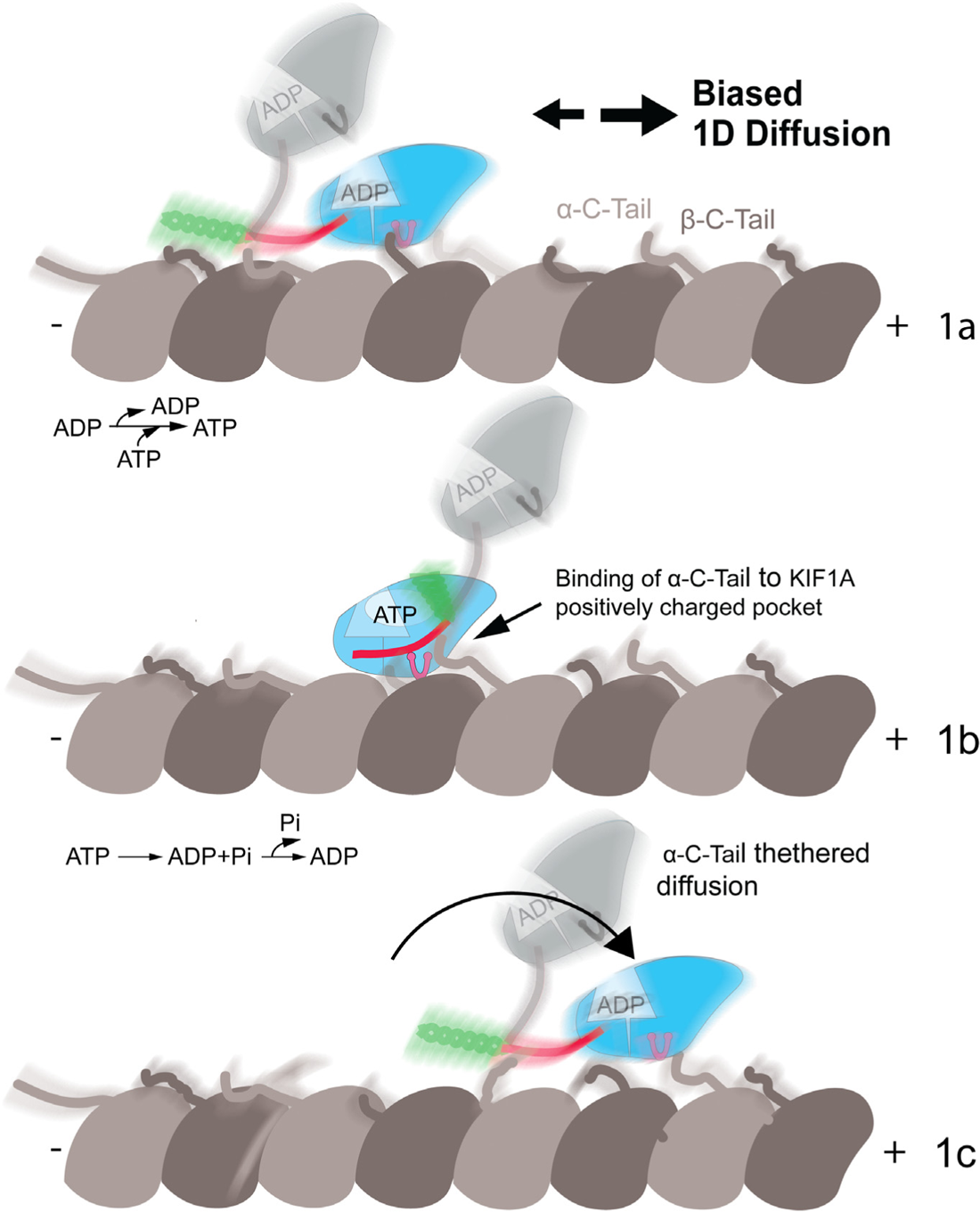
Biased diffusion model. The figure shows a hypothetical model for KIF1A’s MT-plusend-biased one-dimensional-diffusion, inferred from the cryo-EM KIF1A-MT complex structures. In the ADP state, electrostatic interactions between the KIF1A K-loop and the tubulin C terminal tails (Fig. 5b) allow KIF1A to weakly attach to the MT, enabling unbiased diffusion along the MT (step 1a). In this state, one or both KIF1A heads may weakly interact with the MT (a single-head interaction is shown in this figure). Strong attachment of one motor head triggers nucleotide pocket opening and ADP release. Subsequently, ATP binding to the open nucleotide binding pocket induces neck-linker docking (step 1b). In this step, the docked neck-linker, together with the K-loop and part of the coiled-coil domain, creates a pocket of positively charged residues (Fig. 5b). This pocket promotes interaction with the C-tail of α-tubulin, which is located at the MT plus-end relative to the bound motor domain. Completing one ATP-hydrolysis cycle and returning to the ADP state cause the MT-attached head to exit the strongly-bound state, and the neck-linker to undock. Undocking of the neck-linker and the weak attachment to the α-tubulin C-tails results in a step in the MT-plus-direction (step 1c). Combining unbiased one-dimensional diffusion (1b) with MT-plus-end-directed steps (1b-c) results in MT-plus-end-biased one dimensional-diffusion. The proposed model can operate with the single head of a KIF1A monomer. A dimeric KIF1A molecule would have the ability to combine hand-over-hand motion (Fig. 7) with MT-plus-end-biased motility, minimizing the chances of MT detachment during runs and contributing to the characteristic high processivity of KIF1A.

**Supplementary Fig 9.**
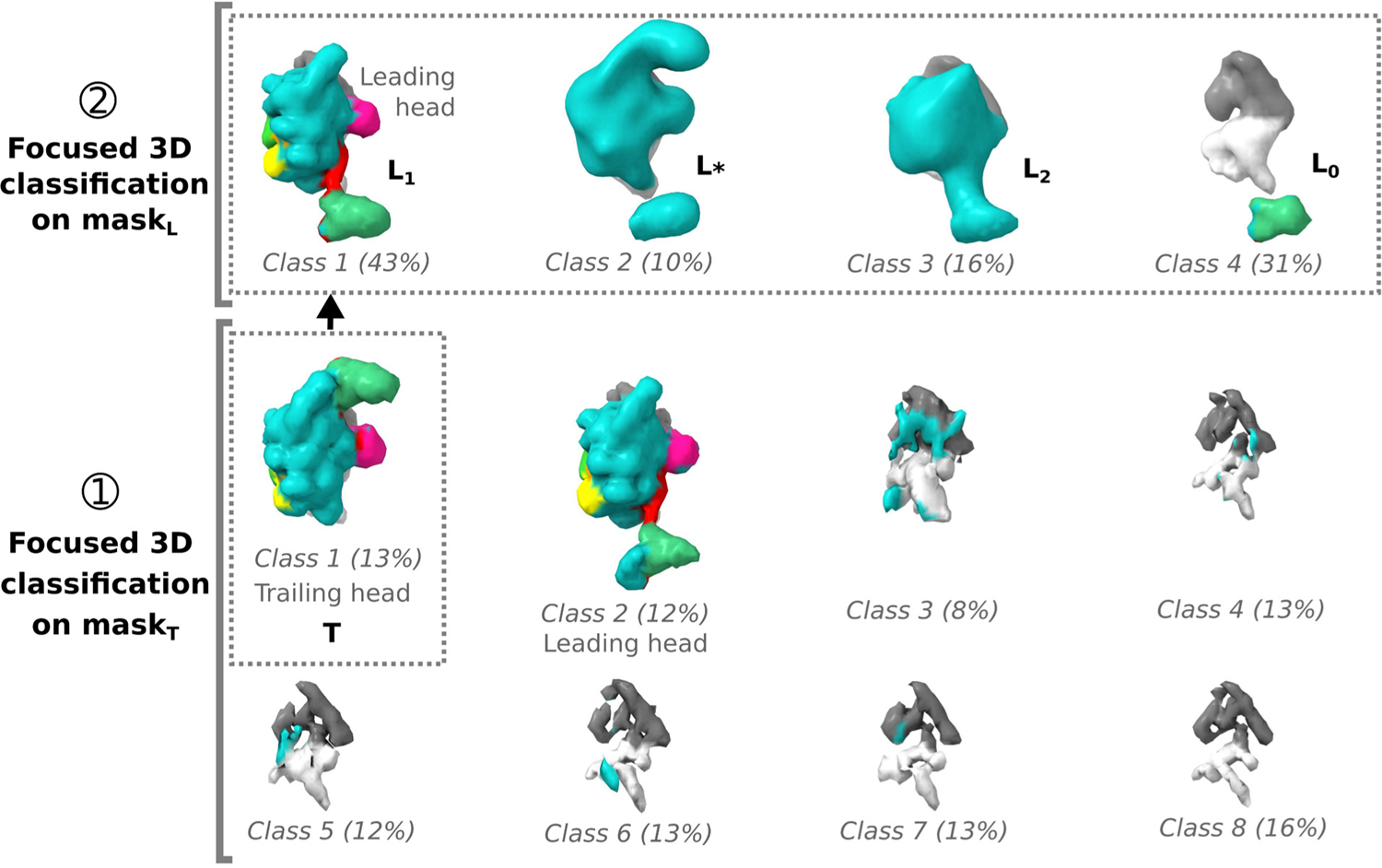
3D classification strategy used for the MT-KIF1A^P305L^-ANP dataset. The same 3D classification strategy employed for the MT-KIF1A-ANP dataset was applied to the MT-KIF1A^P305L^-ANP dataset. The first 3D classification in eight classes focusing on mask_T_ (step ➀) led to the eight class averages displayed in the lower part of the figure. Class 1 corresponds to kinesin motors with the coiled-coil density detected toward the (+) end of the motor (forward), while class 2 corresponds to a leading head with the undocked neck-linker connected to a coiled-coil present toward the (-) end of the motor domain (backward). The other six classes appear undecorated by a kinesin. The single class with a closed conformation was named ‘class T’ and following the same strategy as in the MT-KIF1A-ANP dataset, it was further classified in step ➁ in four classes on the site L (with mask_L_). The corresponding four class averages are displayed in the upper part of the figure. The major class (class 1) corresponds to a motor in the leading head conformation with an undocked neck-linker pulled backward and connected to a coiled-coil whose density is visible. The class TL_1_ therefore corresponds to a two-heads-bound kinesin configuration. Classes 2 and 3, observed at a lower resolution, display similarities with a trailing and leading head, respectively. These were named L_*_ and L_2_. However, due to their lower resolution these classes could not be confidently assigned to these specific states, unlike class L_1_, which exhibited distinct features. Class 4 is devoid of kinesin associated densities so the corresponding motor domains on the site T (class TL_0_) correspond to a one-head-bound configuration, with no leading head bound in the forward direction.

**Supplementary Fig. 10.**
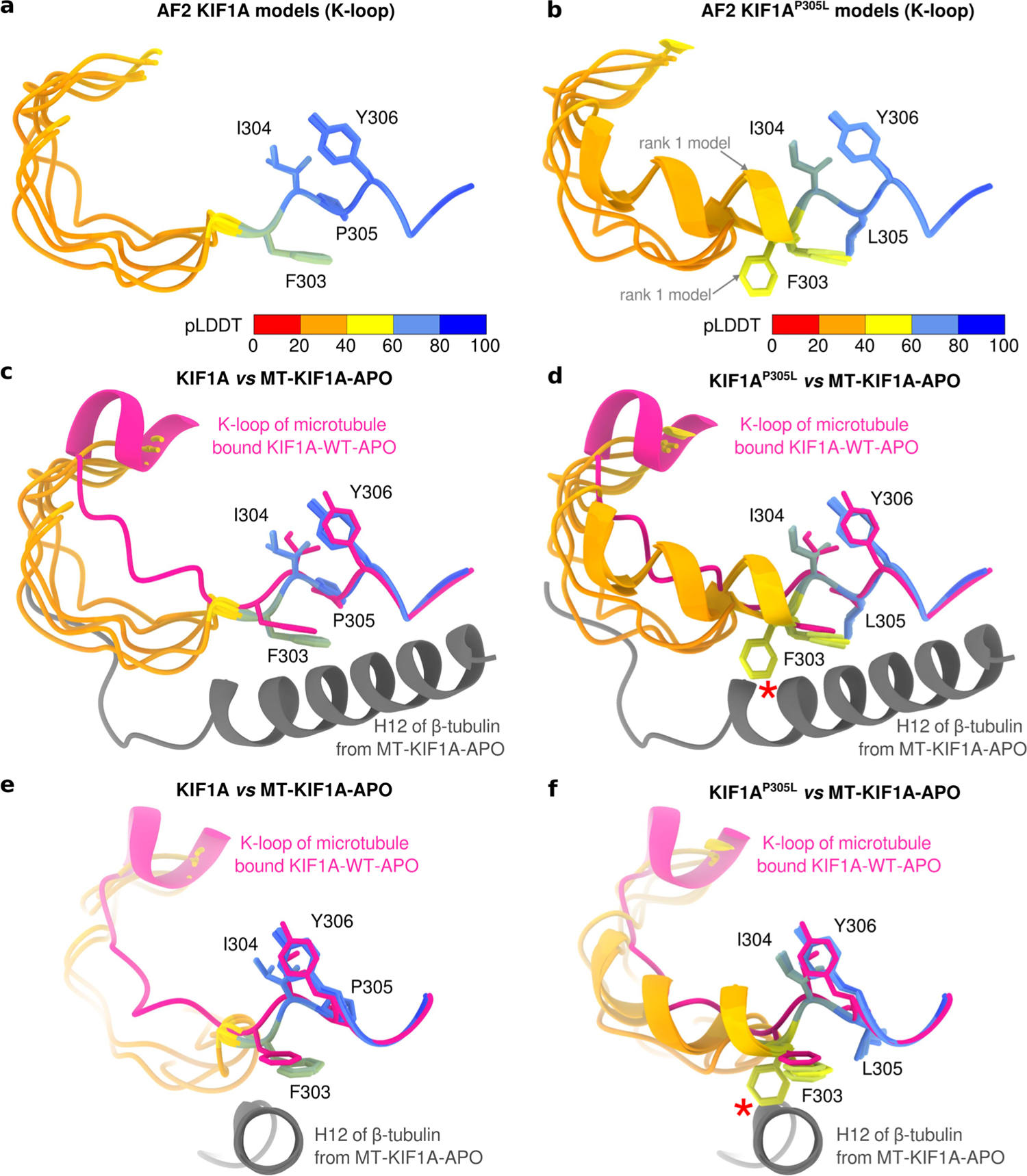
Comparison of AlphaFold2 models of WT KIF1A vs. KIF1A^P305L^ K-loops when not interacting with the MT. **a:** Overlay of the 5 best models of the K-loops from a WT KIF1A motor, generated using the ColabFold^85^ implementation of AlphaFold2 (AF2)^86^. The models were aligned based on residues 304 to 310. Residues are color-coded according to their confidence scores (pLDDT). Note that the structural models of the K-loop are very similar. **b:** Same as (**a**), but for the KIF1A P305L mutant. Notably, the models exhibit greater variability compared to those of WT KIF1A in (**a**). The highest-ranked model is indicated by two arrows. This model together with another model, features a completely different configuration of F303 relative to WT as well as differences in secondary structure in the lower confidence regions of the models that include the more mobile portion of the K-loop containing lysines. **c-d:** Overlays of the same structures as in (**a**) and (**b**) compared with the experimental MT-KIF1A-APO structure. The alignment is performed using the same residues as in (**a-b**). These comparisons were made to assess the local structural changes required for MT binding. In the case of the KIF1A P305L mutant, the alternate conformation of F303 mentioned in (**b**) would strongly clash with helix H12 of β-tubulin (indicated by an asterisk), which is unfavorable for MT binding, as observed. **e-f:** The same alignments as in (c-d), viewed along the axis of helix H12 of β-tubulin from the (+) to the (-) end of the MT.

**Supplementary Table 1.**
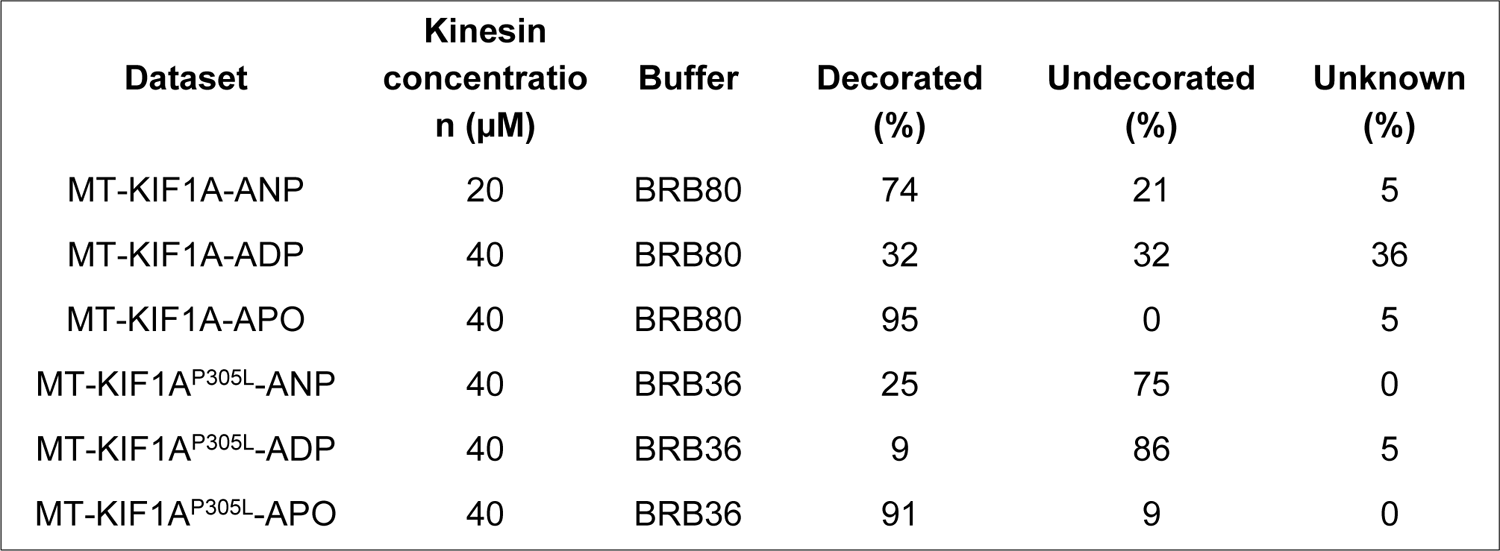
Levels of decoration in each of the cryo-EM datasets. The decorated fraction corresponds to the proportion of the particles images assigned to class(es) for which the class average(s) after the focused 3D classification on the single kinesin site (or site T for ANP datasets) shows a density that could be recognized as being a kinesin motor domain bound to the tubulin dimer. The undecorated fraction corresponds to class averages showing a lack of kinesin motor domain present on the tubulin dimer. In most datasets, there are some low-resolution classes (like Class 6 in Supplementary Fig. 2) for which the class averages show a density that was not reliably assigned as decorated or undecorated and such cases are listed as unknown in the table.

| **Dataset** | **Kinesin**<br>**concentration (μM)** | **Buffer** | **Decorated**<br>**(%)** | **Undecorated**<br>**(%)** | **Unknown**<br>**(%)** |
| --- | --- | --- | --- | --- | --- |
| MT-KIF1A-ANP | 20 | BRB80 | 74 | 21 | 5 |
| MT-KIF1A-ADP | 40 | BRB80 | 32 | 32 | 36 |
| MT-KIF1A-APO | 40 | BRB80 | 95 | 0 | 5 |
| MT-KIF1A<sup>P305L</sup>-ANP | 40 | BRB36 | 25 | 75 | 0 |
| MT-KIF1A<sup>P305L</sup>-ADP | 40 | BRB36 | 9 | 86 | 5 |
| MT-KIF1A<sup>P305L</sup>-APO | 40 | BRB36 | 91 | 9 | 0 |

**Supplementary Table 2.**
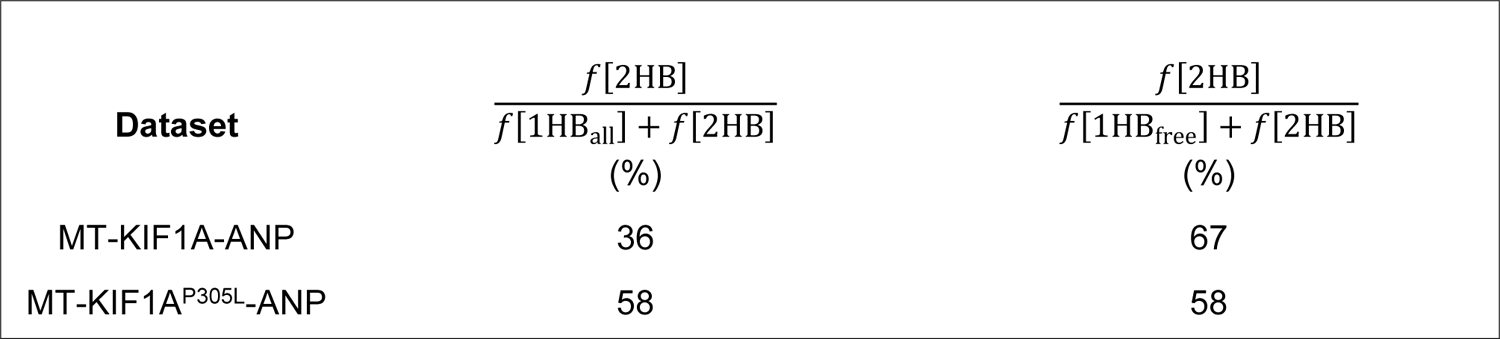
Comparison of the frequencies of one-head-bound and two-heads-bound configurations in the MT-KIF1A-ANP and MT-KIF1A^P305L^-ANP datasets. Due to molecular crowding on the MT, two ratios are given, the right-most one aims to partially account for the large decoration differences between these two datasets (74% for MT-KIF1A-ANP and 25% for MT-KIF1A^P305L^-ANP, see Supplementary Table 1). These ratios use the observed two-heads-bound (2HB) configuration frequency *f*[2HB], the one-head-bound (1HB) configuration frequency *f* [1HB_all_] (corresponding to all one-head-bound configurations detected), and the one-head-bound configuration unconstrained frequency *f*[1HB_free_] (corresponding to particles for which an empty site is present at site L (see Supplementary Figs. 2 and 3). The class frequencies for these two datasets are provided in Supplementary Figs. 2 and 9. Classes with associated class averages at low resolution and/or for which the one-head-bound or two-heads-bound configuration status is unclear are not included in these estimates. For the MT-KIF1A-ANP dataset, the class frequencies combined to estimate the frequency of all the one-head-bound configurations (1HB_all_) are T_1_L_*_, T_1_L_0_, T_2_L_*_, T_2_L_0_, T_3_L_*_, T_3_L_0_; for the one-head-bound configuration unconstrained (1HB_free_) these classes are T_1_L_0_, T_2_L_0_ and T_3_L_0_; and for the two-heads-bound configuration these classes are T_2_L_1_ and T_3_L_1_. For the MT-KIF1A^P305L^-ANP dataset, the class used to estimate the frequency of all the one-head-bound configurations is 1HB_all_, the one used for the one-head-bound configuration unconstrained (1HB_free_) is TL_0_, and for two-heads-bound configuration the class used is TL_1_.

## Notes

### Competing Interest Statement

The authors have declared no competing interest.

### Summary of Updates

Main Figures, Supplementary Figures and parts of the text have been updated to incorporate new data and analysis.

